# RNA-binding protein FXR1 drives cMYC translation by mRNA circularization through eIF4F recruitment in ovarian cancer

**DOI:** 10.1101/2020.07.19.210252

**Authors:** Jasmine George, Yongsheng Li, Deepak Parashar, Shirng-Wern Tsaih, Prachi Gupta, Anjali Geethadevi, Ishaque P. Kadembari, Chandrima Ghosh, Yunguang Sun, Ramani Ramchandran, Janet S. Rader, Hallgeir Rui, Madhusudan Dey, Sunila Pradeep, Pradeep Chaluvally-Raghavan

## Abstract

**Background:** The RNA-binding protein FXR1 (fragile X-related protein 1) has been implicated as an important regulator of post-transcriptional changes of mRNAs. However, its role in mRNA circularization and recruitment of eukaryotic translation initiation factors for protein translation remains obscure. Here, we aimed to investigate the molecular mechanisms and potential clinical applications of FXR1 in ovarian cancer growth and progression.

**Methods:** FXR1 copy number variation, mRNA expression, protein levels, and their association with prognosis were determined in clinical datasets. An orthotopic ovarian cancer model and bioluminescence imaging were used for preclinical evaluation of FXR1 *in vivo*. Reverse phase protein arrays (RPPA) and qPCR arrays were performed to identify FXR1’s key targets and downstream effects. SUnSET and polysome profiling were used to determine the translational effects of FXR1. Immunoprecipitation and immunofluorescence were performed to identify the interaction between FXR1 and cMYC mRNA and eIF4F complex. RNA-immunoprecipitation (RIP), RNA electrophoretic mobility shift assays (REMSA), proximity ligation assays (PLA), and biochemical assays were used to identify the specific site on cMYC mRNA to which FXR1 binds to promote mRNA circularization and translation.

**Results:** We found that amplification and copy-gain of FXR1 increased the expression of *FXR1* mRNA and FXR1 protein in ovarian cancer patients, and these events associated with poor prognosis. We demonstrated that FXR1 binds to AU-rich elements (ARE) within the 3’ untranslated region (3’UTR) of cMYC. As a consequence, FXR1 binding to cMYC 3’UTR leads to the circularization of mRNA and facilitated the recruitment of eukaryotic translation initiation factors (eIFs) to translation start site for improving protein synthesis.

**Conclusion:** We found that FXR1 upregulates a known oncogene, cMYC, by binding to AU-rich elements within the 3’UTR, leading to the recruitment of the eIF4F complex for cMYC translation. Our findings uncover a novel mechanism of action of FXR1 in tumorigenesis and provides opportunities to use FXR1 and its downstream effectors as biomarkers or therapeutic targets in ovarian and other cancers.

## Introduction

Copy number variations (CNVs), such as genomic amplification, copy number gain, or deletion, are frequent events in ovarian cancer [1]. Among CNVs, amplification of the 3q26 locus is seen in ∼30% of high-grade serous ovarian cancers (HGSOC). Previously, it was demonstrated that many genes, such as *PI3KCA* [2], *EVI1* [3], and *TERC* [4], are parts of the 3q26 locus that contribute to the development and progression of ovarian and other cancers. Recently, our group reported that non-coding RNAs, such as the microRNAs miR569 and miR551b, that are amplified as part of the 3q26.2 locus contribute to the oncogenesis and progression of breast and ovarian cancers [5–7]. Collectively, these studies suggest that 3q26 is important in ovarian and other cancers because many genes in this amplicon promote oncogenesis either individually or by cooperating with other genes in this locus or with their downstream targets or actions.

By analyzing high-resolution single nucleotide polymorphism (SNP) array data of TCGA, we determined that FXR1 (fragile X-related protein 1), which is located in the 3q26.3 chromosomal locus, is highly amplified or copy-gained in ovarian cancer and many other cancers, including squamous cell carcinoma of the lung, cervix, head, and neck. FXR1, which is highly conserved in mammals [8], is a member of the fragile X-related (FXR) family of RNA-binding proteins (RBPs). The major biological functions of RBPs include regulation of RNA stability, splicing, and translation [9]. Recent studies have shown that FXR1 is a key promoter of tumor progression, which is critical for the growth of many cancers such as non-small cell lung cancer (NSCLC) [10], prostate cancer [11], glioma [12], and oral squamous cell carcinoma (OSCC) [13]. It has been reported that FXR1 binds to AU-rich elements (AREs) within the 3′untranslated region (3′UTR) and enhances the stability of *TNF-*α and *COX2* mRNAs [14, 15]. AREs, which are located within the 3 ′UTRs of mRNAs, are key target sequences of RBPs, and RBPs influence mRNA stability, translation progress, and alternative pre-mRNA processing [16]. The conserved nucleotide sequence motif in AREs is AUUUA, which occurs in variable length repetitions in the 3′ UTR of mRNAs [17]. However, it is unclear how FXR1 stabilizes its target MRNAs and promotes their translation and whether it is important for the pathophysiology and progression of ovarian cancer. In this paper, we describe how FXR1 binds to the ARE within *cMYC* mRNA, enhancing its translation to cMYC oncoprotein and a multifunctional transcription factor, which is important for the growth and aggressiveness of ovarian cancer.

## Results

### FXR1 copy number amplification associates with high expression of FXR1 and poor cancer outcomes

The 3q26 amplicon is large and structurally complex, consisting of about 50 genes and non-coding elements [7, 18]. Studies by us and others have demonstrated that multiple genes and non-coding RNAs in this amplicon contribute to both tumor initiation and progression either alone or cooperatively [6, 19–22]. To identify genes within the 3q26 locus that contribute to ovarian cancer oncogenesis, we used high-resolution SNP-based copy number analysis of 579 HGSOC and 1080 breast cancers in the Cancer Genome Atlas (TCGA). Our analysis revealed that FXR1 is highly amplified or copy-number gained in >40% of HGSOC patients and >25% of breast cancer patients (**Fig 1A and S1A**). To investigate whether FXR1’s expression profile is also altered in other cancers due to CNV, we interrogated TCGA dataset cBioPortal (https://www.cbioportal.org/) and Clinical Proteomic Tumor Analysis Consortium (CPTAC) for CNV, gene expression, and protein changes. We also interrogated UALCAN (http://ualcan.path.uab.edu/) for alterations in FXR1 expression in a variety of human cancer samples. We found that FXR1 is highly amplified in several human malignancies, particularly lung, ovarian, cervical, colon, and breast cancers **(Fig 1B and S1B)**. Strikingly, FXR1 CNVs associated with increased expression and levels of both *FXR1* mRNA and *FXR1* protein in ovarian cancer patients in the TCGA dataset (**Fig 1C**). Similarly, FXR1 CNV associated with *FXR1* mRNA in breast cancer, lung adenocarcinoma (LUAD), and lung squamous carcinoma (LUSC) patients in that dataset (**Fig S1C**). Taken together, our data suggest that a gain in copy number or amplification of the FXR1 gene leads to increased expression of *FXR1* mRNA and, consequently, to higher levels of FXR1 protein. Next, we sought to determine whether *FXR1* mRNA associated with patient outcomes in a publicly available ovarian cancer dataset [23]. We found that high *FXR1* mRNA expression associated with worse overall and recurrence-free survival in those patients (**Fig 1D**).

**Fig 1.**
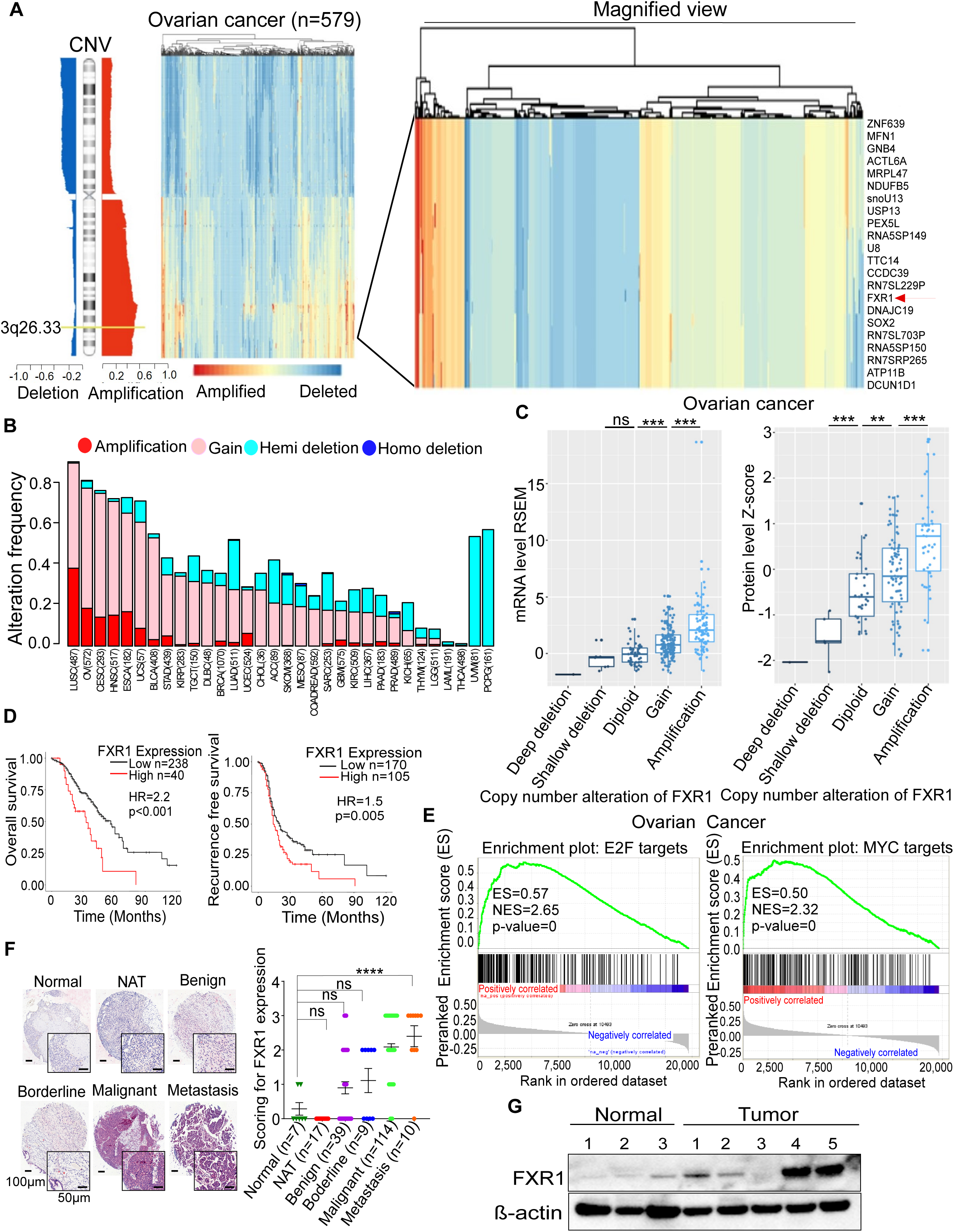
*FXR1* mRNA and FXR1 protein associated with the progression of ovarian cancer. **(A)** Heatmap of chromosome 3q26.3 locus amplification in TCGA ovarian cancer patient dataset (n=579). Representative genes in this amplicon is magnified, where red arrow indicate position of FXR1. **(B)** Frequency of FXR1 alterations across human cancers in TCGA database. **(C)** Box plots show the association of copy number alterations of FXR1 with *FXR1* mRNA (left panel) and FXR1 protein (right panel). These considered with respect to the median expression of all data in the TCGA ovarian cancer data set. **p<0.01, ***p<0.001 compared between indicated groups. ns: non-significant. p-value was determined by two-sided Wilcoxon rank sum test. **(D)** Kaplan– Meier plot shows FXR1 expression-based outcome of overall survival and recurrence-free survival in 278 ovarian cancer patients in the Tothil et al. (Bowtell) dataset. Patients were stratified according to the median expression of FXR1 and p-value was determined by log-rank test. **(E)** GSEA analysis demonstrates the enrichment score of indicated functional annotation marks based on the correlation between expression of all genes and FXR1 expression in the TCGA ovarian cancer samples. ES: enrichment score, NES: normalized enrichment score. **(F)** Immunohistochemistry (IHC) analysis of ovarian cancer tissue microarray was performed for FXR1. Vector Red chromagen staining was performed and IHC scoring was performed based on the positive staining of FXR1 (red color). Sample numbers are included as normal (n=7), normal adjacent tissue (NAT, n=17), benign (n=39), borderline (n=9), malignant (n=114) and metastasis (n=10) samples from two different ovarian cancer tissue microarrays (Fig S1E) are shown. Individual tissue cores were blindly scored for FXR1 staining intensity as negative (0), trace, weak (1+), moderate (2+), or strong (3+) staining in 50% of the cells examined. Error bars indicate SEM. ****p<0.0001 compared to normal tissues by Anova test. ns: non-significant. **(G)** Western blot analysis of FXR1 levels in normal and ovarian cancer patients’ tissue. β-actin was used as a loading control.

Next, we employed Gene Set Enrichment Analysis (GSEA) to identify potential functional gene sets that associated with high expression of FXR1. First, we ranked all the protein-coding genes based on their correlation with FXR1 expression; then we focused on functional gene sets related to cancer hallmarks. Importantly, this GSEA analysis found that high expression of FXR1 showed the greatest association with two enrichment annotations that included MYC targets and E2F1 targets in TCGA ovarian, breast, LUAD, and LUSC datasets (**Fig 1E and S1D**).

To confirm the data from our bioinformatic analyses, we quantified FXR1 protein in ovarian cancer tissues by immunohistochemical (IHC) analysis, using two tissue microarrays (TMAs) that included 172 ovarian cancer samples, 17 adjacent normal tissues, and 7 normal tissues (**Fig S1E and Table S1, Table S2**). We quantified FXR1 protein levels by scoring positive staining intensity in each tissue spot in the array.

Importantly, malignant and metastatic samples displayed ∼3-fold higher expression of FXR1 compared to normal and benign tissues, with the metastatic tissues showing the highest levels (**Fig 1F**). In contrast, immunoblotting of normal human ovarian lysates detected barely any FXR1 (**Fig. 1G**); whereas we found that FXR1 protein was expressed at high levels in 3 of 5 cancer tissues and at low to modest levels in the other two (**Fig 1G**). Thus, FXR1 was highly expressed in most of the HGSOC cells except OVCAR3, which expressed only low levels **(Fig S1F and S1G)**. Taken together, our data suggest that FXR1 is highly expressed in a large subset of ovarian cancer patients due to CNV. Such aberrant overexpression of FXR1 in ovarian cancer cells was confirmed by quantitative PCR and immunoblotting. Collectively, these results suggest that FXR1 likely plays a critical role in ovarian cancer progression.

### Depletion of FXR1 reduces oncogenic properties of ovarian cancer cells

To determine the effects of FXR1 in ovarian cancer cell lines, we used two different siRNA sequences that specifically target and inhibit FXR1 **(Fig 2A).** OVCAR5 and HeyA8 cells transfected with siFXR1 reduced their proliferation and formed fewer colonies than cells transfected with control siRNA **(Fig 2B and 2C)**. We also observed that loss of FXR1 reduced the invasiveness of ovarian cancer cells 24h after the cells were seeded onto Matrigel-coated trans-well inserts (**Fig 2D**). These data suggest that ovarian cancer cells need to express FXR1 to proliferate and form colonies.

**Fig 2:**
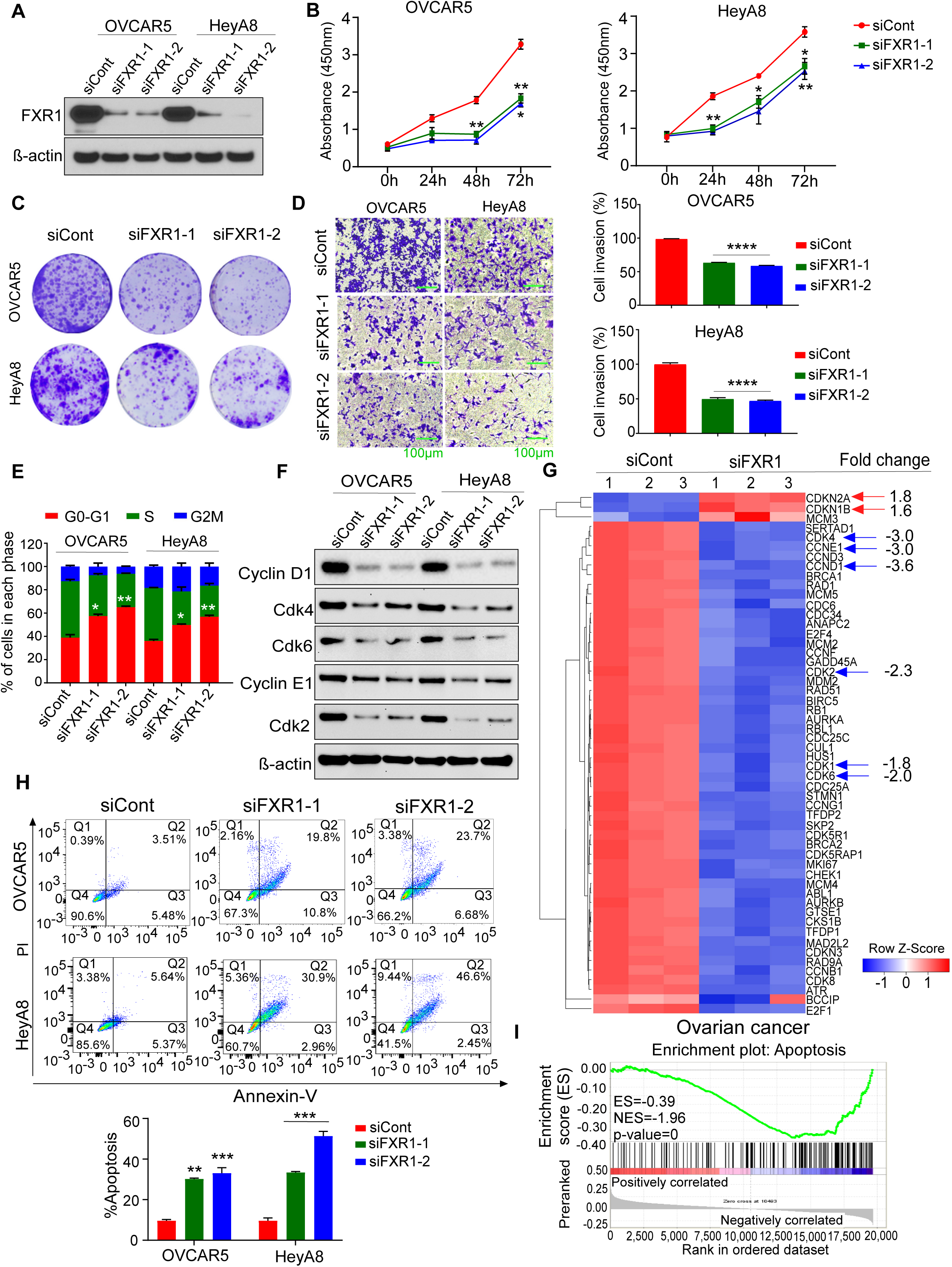
FXR1 knockdown inhibited the growth of ovarian cancer cells by promoting the apoptosis. **(A)** OVCAR5 and HeyA8 cells were transfected with two different siRNAs targets FXR1 (siFXR1-1 and siFXR1-2) or control siRNAs (siCont) and Western blot was performed 48h after transfection using antibodies indicated. β-actin was used as a loading control. **(B)** OVCAR5 and HeyA8 cells were transfected using the siRNAs and cell survival rate was determined using the CCK-8 assay at indicated time points < 0.05, **p<0.01 compared to siCont by (n=5). Results are shown as ± SEM. *p < 0.05, **p<0.01 compared to siCont by Student’s t-test. OVCAR5 and HeyA8 cells were transfected using the siRNAs as in **A** and subjected to **(C)** colony formation assay and **(D)** invasion assay, results are shown as mean of three technical replicates ± SEM. ****p<0.0001 compared to siCont by Student’s t-test. **(E)** Flow cytometry analysis of cell cycle of ovarian cancer cells transfected with siRNAs. Results are shown as mean of three technical replicates ± SEM. *p < 0.05, **p<0.01 compared to siCont by Student’s t-test. **(F)** Western blots of the lysates from cells in **E** were performed using the antibodies indicated. β actin was used as a loading control. **(G)** qPCR array was performed using the cDNA prepared from the HeyA8 cells were transfected with siCont or siFXR1. Gene expression is represented as Log2 of Ct values and with p>0.05 compared to siCont. Genes that were ≥ 1.5-fold down (blue color)- or upregulated (red color) by FXR1 were included in the heat map (n=54). **(H)** Cells transfected with siCont or FXR1 siRNAs were double-stained with annexin V-FITC and PI, and then subjected to flow cytometric analysis after 48h of transfection (top). Results were presented as percentage of apoptotic cells in both siCont and FXR1 siRNAs treated cells and shown as mean of three technical replicates ± SEM. **p<0.01, ***p<0.001 compared to siCont by Student’s t-test (bottom). **(I)** GSEA analysis demonstrates the enrichment score of indicated functional annotation marks based on FXR1 expression in the TCGA ovarian cancer samples. ES: enrichment score, NES: normalized enrichment score.

Next, we performed cell cycle analysis to determine FXR1’s effects on cell cycle phases. Loss of FXR1 increased the number of cells in the G0/G1 phase with a concomitant decrease in the number of cells in the S and G2/M phases (**Fig 2E and S2A**). To corroborate these effects, we determined levels of cyclins and cyclin dependent kinases (CDKs), which are important for the G1 growth phase. Loss of FXR1 reduced levels of Cyclin D1, Cyclin E1, CDK4, CDK6, and CDK2 (**Fig 2F**). Next, we used a qPCR array to determine the expression of genes associated with cell cycle regulation, cell survival, and proliferation (**Table S3**). Many of the targets related to the cell cycle, survival, and proliferation. For example, expression of *CDK4, CCNE1, CCND1, CDK2, CDK1,* and *CDK6* in HeyA8 ovarian cancer cells decreased remarkably upon loss of expression of FXR1, whereas expression of CDK inhibitors such as *CDKN2A* (codes for INK4A and ARF) and *CDKN1B* (p27) increased (**Fig 2G**).

Consistent with these changes, we also observed prominent morphological changes associated with cell death (such as nuclear condensation and DNA fragmentation) when FXR1 was knocked down (**Fig S2B**). To investigate apoptosis in the knocked-down cells, we performed flow cytometry, using Annexin V-FITC, which binds to phosphatidyl serine flipped outside on the cell membrane when a cell undergoes apoptosis. In this analysis, we found that the depletion of FXR1 increased the apoptosis of both OVCAR5 and HeyA8 cells as plotted as Q2 populations (**Fig 2H)**. Next, we performed GSEA, and found that *FXR1* mRNA levels inversely correlated with the functional annotation mark of apoptosis based on the correlation of genes with *FXR1* mRNA in the TCGA datasets of ovarian cancer, breast cancer, and LUSC (**Fig 2I and S2C**).

We also found that loss of FXR1 increased cleaved caspase-3 and caspase-7 activities, which are surrogates for apoptosis in ovarian cancer cells (**Fig S2D**). To further determine changes in levels of proteins related to cell death, we performed a protein array with apoptotic markers, finding that FXR1 depletion increased levels of the pro-apoptotic proteins BAX, cytochrome-C, death receptors FADD, FAS, p21, p27, and phospho-p53 (S15) and decreased levels of pro-survival proteins such as HSP-60 and SURVIVIN (**Fig S2E**). In conjunction, our immunoblot also showed that FXR1 deficiency upregulated levels of BAX, P27, and P21 and reduced levels of BCL2 (**Fig S2F**). Together, our results suggest that FXR1 knockdown inhibits cell growth, prevents cell cycle progression, and activates cell death pathways in cancer cells.

### FXR1 regulates the expression of cMYC in ovarian cancer cells

Many RBPs are known to regulate the stability of mRNAs and to modulate protein translation [24]. Therefore, we determined whether FXR1 contributed to global or gene-specific translational regulation in HeyA8 ovarian cancer cells. Here, we performed a surface sensing of translation (SUnSET) assay, using puromycin, a structural analog of aminoacyl tRNAs (specifically tyrosyl-tRNA; **Fig S3A**). During elongation of a nascent polypeptide chain, the binding of puromycin prevents a new peptide bond from being formed with the next aminoacyl-tRNA [25] (**Fig 3A and S3A**). Therefore, puromycin terminates peptide elongation, leading to the release of a truncated puromycin-bound peptide from the ribosome.

**Fig 3:**
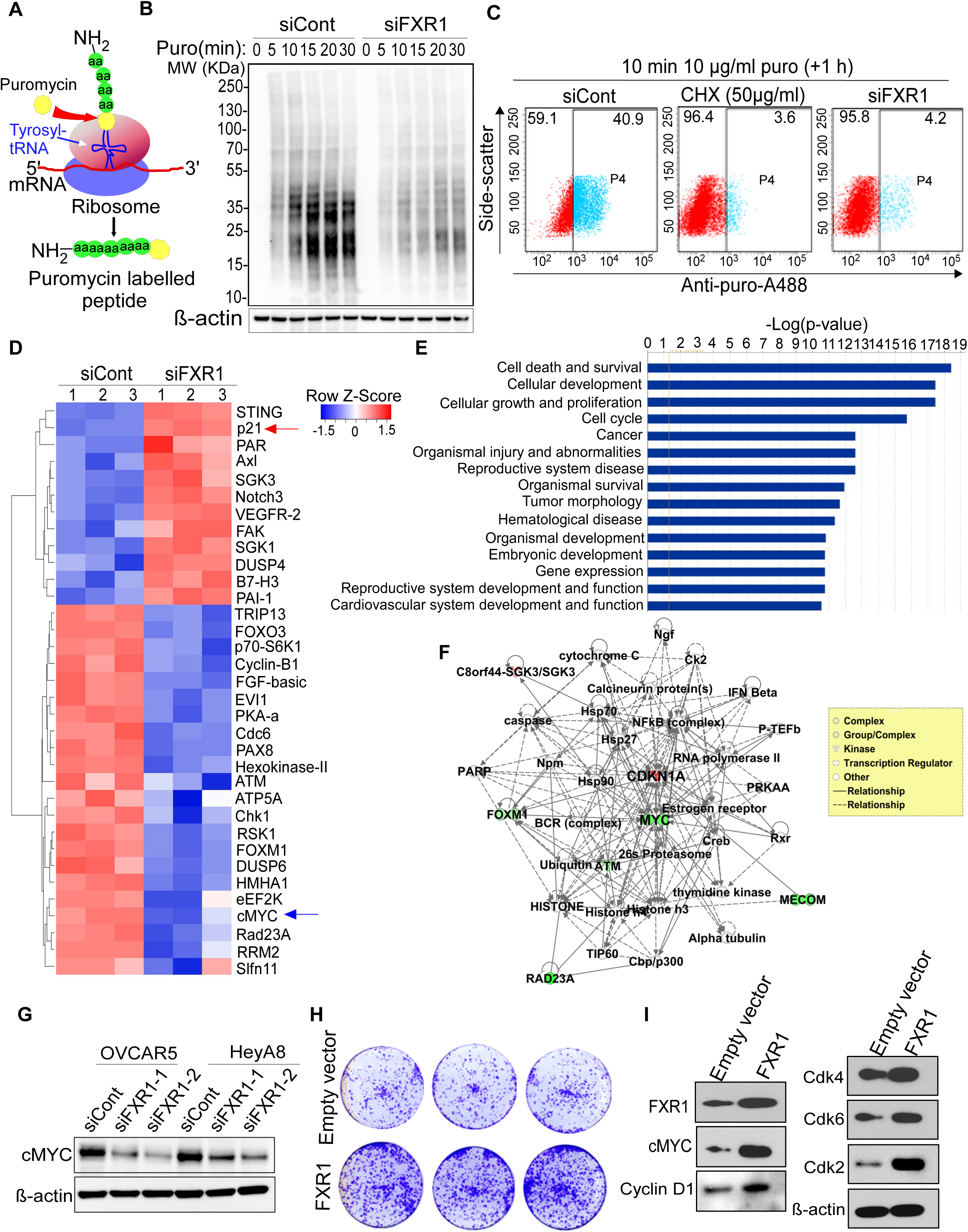
FXR1 alters the overall protein translation and promotes the levels of cMYC protein. **(A)** Schema illustrates the principle of surface sensing of translation (SUnSET) assay how puromycin acts as a structural analog of tyrosyl-tRNA incorporated in a growing peptide chain. **(B)** Western blot analysis of proteins labeled with puromycin (puro) using anti-puromycin antibody in time-dependent manner as indicated in HeyA8 cells treated with either siCont or siFXR1. β-actin was used as a loading control. **(C)** β-actin was used FACS sorting of the puromycin labelled cells using the anti-puromycin antibody tagged with Alexa Fluor 647 in the HeyA8 cells were transfected with either siCont or siFXR1 or treated with cycloheximide (CHX). **(D)** Heatmap of the differentially expressed proteins based on Log2 fold change (n=34) identified by Reverse phase protein array (RPPA) analysis with a p-value of < 0.05 compared to siCont. Normalized Log2 values of RPPA signal were used in this analysis. Blue and red values represent down- and upregulated proteins respectively. **(E)** Functional network analysis was performed using the differentially expressed proteins obtained from RPPA (n= 34, Log2 fold change) by Ingenuity Pathway Analysis (IPA). The top 15 Molecular and Cellular Functions were ranked based on p-value, and the bars represent inverse log of the p-value (x-axis). A p-value < 0.05 by Fisher’s exact test was considered to select statistically significant pathway annotation. **(F)** Top gene networks from the pathways generated by the Ingenuity Pathway Analysis (IPA) suite. Green highlighted proteins indicate downregulated and red indicate upregulated upon siFXR1 treatment. Solid lines indicate direct protein–protein interactions, dashed lines indicate potential protein–protein interaction, and arrows imply directionality of protein–protein regulation with CDKN1A (p21) and cMYC (score of 14) as central regulators under the function ‘Cell cycle, cellular development and tissue development function’. Genes are represented by nodes with their shape representing the type of molecule/functional class, and the relationship between the nodes are indicated by edges. **(G)** Ovarian cancer cells were transfected with indicated siRNAs and immunoblot analysis was performed using the lysates prepared after 48h of transfection using indicated antibodies. β-actin was used as a loading control. **(H)** Colony formation assay for OVCAR3 cells were transfected with FXR1 expressing vector (pReceiver-M39-FXR1-FLAG) or control empty vector (pReceiver-M39-FLAG). **(I)** Western blots of the lysates from cells in **H** were performed using the antibodies indicated. β-actin was used as a loading control.

HeyA8 cells were transfected with control siRNA or FXR1 siRNA, treated with a low dose of puromycin (10 µg/ml) at the indicated time points up to 30 minutes and immunoblotted with anti-puromycin antibody. Our SUnSET assay demonstrated that inhibition of FXR1 reduced the overall translation of proteins in a time-dependent manner (**Fig 3B**). To complement this result, we quantified the puromycin-labeled cells from the SUnSET assay by using fluorescently activated cell sorting (FACS). We found that knocking down FXR1 reduced overall protein synthesis, as happened similar when the cells were treated with the translational inhibitor cycloheximide (CHX) (**Fig 3C**).

These data motivated us to identify the key proteins that FXR1 regulates. We used a reverse phase protein array (RPPA)—a functional proteomic screening assay—with the HeyA8 cells transfected with control siRNA or FXR1 siRNA. This high-throughput antibody-based technique was developed to evaluate proteins known to associate with various signaling networks in cancer cells [26]. In our RPPA assay with knocked-down FXR1, we found that 34 proteins were altered significantly compared to the control siRNA-transfected cells. Among the 34, 12 were upregulated and 22 were downregulated (**Fig 3D and S3B**). Consistent with our finding that FXR1 inhibited apoptosis and promoted oncogenesis (see **Fig 2, Fig 4** and **Fig 5**), we observed that silencing FXR1 reduced levels of many proteins known for their oncogenic functions, including cMYC, EVI1, CHK1, FOXM1, Cyclin B1, and CDC6. Notably, we also found that loss of FXR1 upregulated levels of CDKN1A (p21), DUSP4, FAK1, and PAI1, which are mainly involved in the cell death mechanisms of cancer cells (**Fig 3D**).

**Fig 4:**
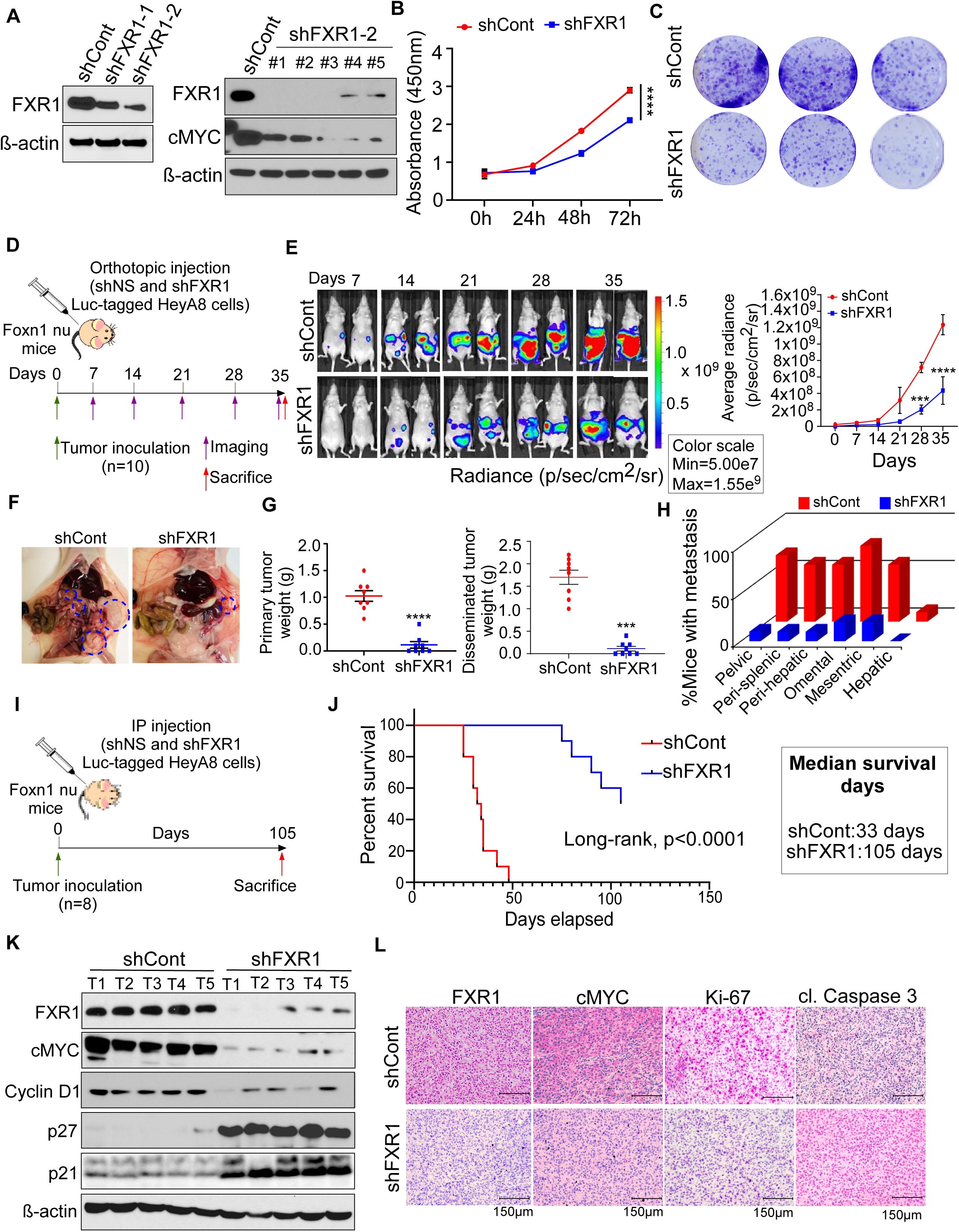
Stable knockdown of FXR1 reduced the overall tumor growth, metastasis and prolonged survival of ovarian xenograft mouse model. **(A)** Western blot analysis for confirming the knockdown of FXR1 in HeyA8 cells prepared using pLKO.1 vector express two independent shRNAs constructs of FXR1 (1: TRCN0000160812 and 2: TRCN0000160901) or empty control pLKO.1 vector (shCont). β-actin was used as a loading control. **(B)** shRNA hairpin number 2 (shFXR1-2) were used to make single cell derived stable cell colonies in the presence of puromycin (8µg/ml). Lysates were prepared from the colonies (shFXR1 #1-#5) and immunoblotted using the antibodies indicated (right). β-actin was used as a loading control. **(C)** Cell survival rate of shFXR1 and shCont HeyA8 cells were determined using the CCK-8 assay at the indicated time point (n=8). Results are shown as ± SEM. ****p<0.0001 compared to shCont cells by Student’s t-test. **(D)** Stable cells from **B** were plated and the colonies formed were photographed on day 10. **(E)** Timeline of establishment of ovarian cancer xenograft mouse models using 25×10^3^ luciferase tagged HeyA8 cells (shCont and shFXR1) were orthotopically inoculated into the left ovary bursa of female athymic nude mice (n=10/group, green arrow). **(F)** Mice from both groups were imaged using biophotonic IVIS at the indicated time points and representative photographs were presented (left). Line graph indicates the average radiance of signaling intensity. Error bars indicate SEM. ***p<0.001, ****p<0.0001 compared to shCont by two-way Anova test. **(G)** Representative image of a mouse from shCont and shFXR1 groups were surgically opened and photographed. Areas circled in blue indicate tumor growth and metastatic locations. **(H)** Primary and disseminated tumors were collected from **E** and total tumor weight was recorded. Error bars indicate SEM. ***p<0.001, ****p<0.0001 compared to shCont mice by Student’s t-test. **(I)** Bar graph shows the percentage of mice with metastasis in the indicated organs in both shCont and shFXR1 groups. **(J)** Schema shows the plan and timeline we used for survival analysis. 25×10^3^ luciferase tagged HeyA8 cells (shCont or shFXR1) were intraperitoneally (IP) inoculated into female athymic nude mice (n=8/group, green arrow). **(K)** Kaplan Meier analysis was performed using the death record of mice from each group in **I**. p-value was determined by log-rank test. **(L)** Western blot analysis of lysates was collected from tumor tissues selected from each group (n=5) using indicated antibodies. β-actin was used as a loading control. **(M)** Representative images (×20 magnification) of indicated antibodies prepared using the ovarian tumor tissues collected from shCont and shFXR1 mice from **I**.

**Fig 5:**
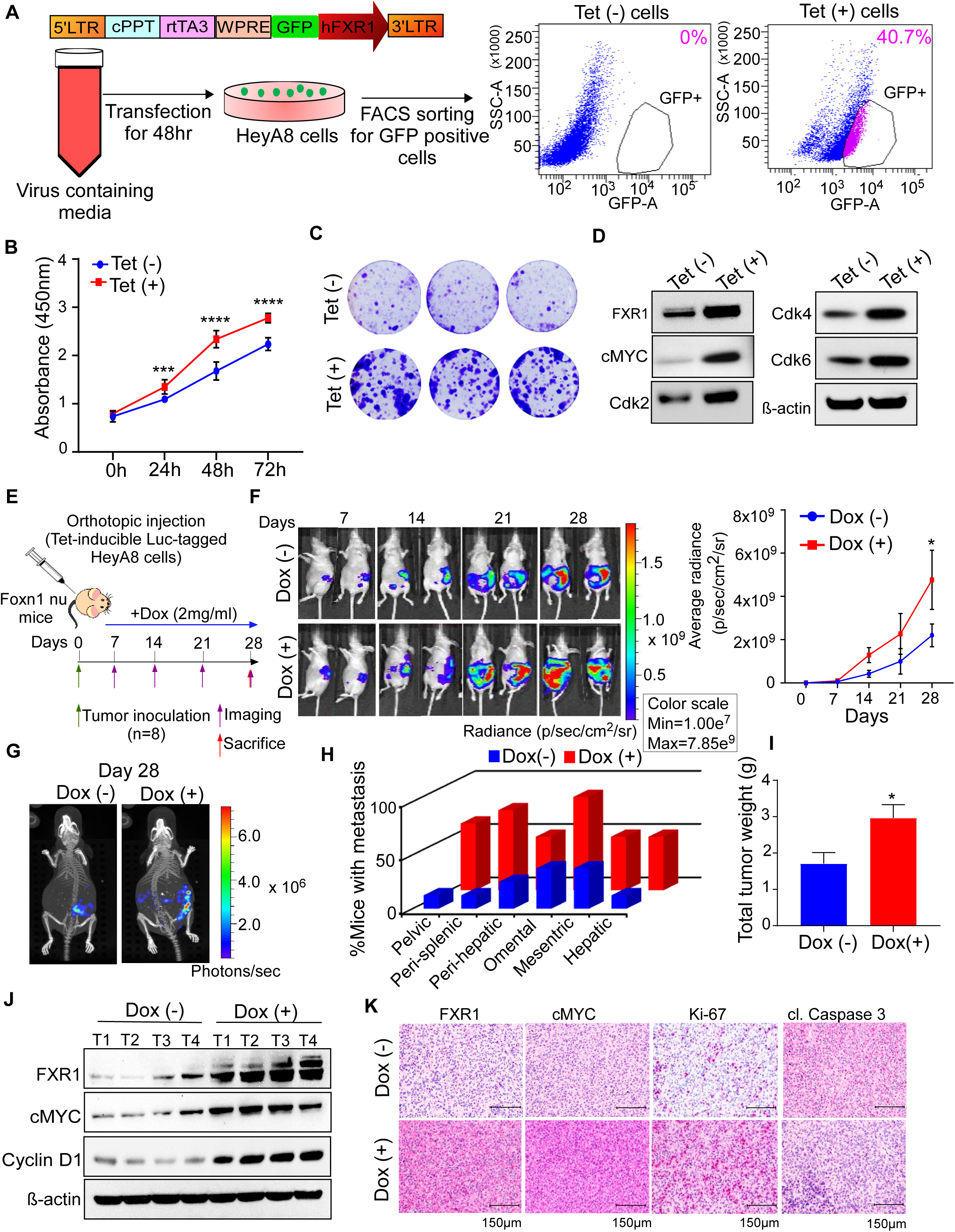
FXR1 overexpression by Tet-inducible system increases tumor growth and metastasis in ovarian xenograft mouse model. **(A)** Vector map of tetracycline inducible pFLCt construct, where we cloned hFXR1-and GFP, which was used for the preparation of hFXR1-and GFP stably expressing HeyA8 cells. Stable cells which express inducible GFP and FXR1 were selected by FACS after treatment with tetracycline (1µg/ml). **(B)** Proliferation of HeyA8 cells which stably express inducible GFP and FXR1 was determined using the CCK-8 assay at the indicated time points (n=10). Results are shown as mean of ± SEM. ***p<0.001, ****p<0.0001 compared to the cells were not treated with tetracycline by Student’s t-test. **(C)** HeyA8 cells from **A** were plated in the presence or absence of tetracycline for 10 days and the colonies were photographed. β-actin was used as a loading control. **(D)** Cell lysates were prepared from **A** and immunoblot was performed using the antibodies indicated. **(E)** Timeline of establishment of ovarian cancer xenograft mouse models using the HeyA8 cells stably express GFP and FXR1. 25×10^3^ tetracycline inducible luciferase tagged HeyA8 cells were orthotopically inoculated into the left ovary bursa of female athymic nude mice. Mice were feed with or without Doxycycline (2mg/ml) in drinking water (n=8/group) and imaged using IVIS imager. **(F)** Mice (n=8/group) were imaged from **E** at the indicated time points and the photographs of representative mice were presented (left) and the intensity of luciferase reporter signals were plotted (right). Error bars indicate SEM, *p < 0.05 compared to Dox (+) group by two-way Anova test. **(G)** Representative 3D PET/CT images of mouse from Dox (-) and Dox (+) group from **F** on the indicated day point. **(H)** Bar graph shows the percentage of mice with metastasis in the mice from **F** at indicated organs. **(I)** Total tumor weight of the tumors isolated from **F** on day 28. Error bars indicate SEM, *p < 0.05 compared to Dox (+) group by Student’s t-test. **(J)** Western blot analysis of lysates was collected from tumor tissues selected from each group (n=4) using indicated antibodies. β-actin was used as a loading control. **(K)** Representative images (×20 magnification) of indicated antibodies prepared using the ovarian tumor tissues from **E**.

Next we determined the proteins that are central driver of the canonical protein networks; by performing the pathway enrichment analysis using Ingenuity Pathway Analysis software (IPA; Ingenuity Systems Inc) using the proteins which were significantly altered in our RPPA. Our analysis identified top-five enriched pathways which are crucial for cancer progression such as cell death and cell survival, cellular development, cellular growth and proliferation, cell cycle and cancer a (**Fig 3E**). We then selected the top three gene networks by ranking functional interaction scores (**Fig 3E and S3C**). This approach identified cMYC as the most downregulated protein, when we knocked down FXR1 as a center of a gene network. We also found that p21 was the most upregulated protein in the identified pathways when FXR1 was depleted (**Fig 3F**). In addition to the above proteins, we identified other protein networks that included Cyclin E, Cyclin D, CHEK1, E2F, RAF, FGF, SGK1, and NOTCH3 (**Fig S3C**).

To confirm the RPPA results, we immunoblotted ovarian cancer cell lysates collected when FXR1 was knocked down. This assay showed that silencing FXR1 remarkably reduced the expression of cMYC protein (**Fig 3G**). To complement these approaches, we overexpressed FXR1, using FXR1 cloned into pReceiver-M39-FLAG or control vector pReceiver-M39 transfected in OVCAR3 cells which express low levels of FXR1 (**Fig S1G**). This assay showed that overexpression of FXR1 increased colony formation and upregulated levels of cMYC, CDK4, CDK6, and CDK2 in the ovarian cancer cells (**Fig 3H and 3I**). Next, we used a tissue microarray (TMA) prepared from ovarian cancer patients to determine the association between FXR1 and cMYC. Based on the IHC scores (see **Fig S1E and S3D**), we found a strong association between levels of cMYC and FXR1 in both benign (r=0.8054, *p*<0.0001) and malignant (r=0.6604, *p*<0.0001) ovarian carcinoma tissues (**Fig S3D**). Taken together, our data demonstrate that FXR1 regulates the expression of many proteins that are important for several key oncogenic features, including cell cycle regulation and proliferation, where cMYC oncoprotein is likely to be the direct target of FXR1.

### Depletion of FXR1 suppresses the growth of ovarian cancer cells and improves the survival of mice bearing orthotopic ovarian tumors

Based on our *in vitro* data, we postulated that low levels of FXR1 reduce tumor burden *in vivo*. To test this hypothesis, we stably knocked down FXR1 in a HeyA8 cell line, using two different short hairpin RNAs (shRNA) (TRCN0000160812 labeled as shFXR1-1 and TRCN0000160901 labeled as shFXR1-2) or a nonspecific control shRNA (shCont) cloned in pLKO.1 vector. Of the two sequences, TRCN0000160901 (shFXR1-2) produced the greatest decrease in FXR1 levels (**Fig 4A**). Stably knocked-down cells and control cells were selected after puromycin treatment (8 µg/ml), which confirmed the decreased FXR1 levels in all the selected single-cell clones. As expected, we found that the level of FXR1 protein was markedly lower in the shFXR1 cells compared with the shCont cells in all the clones (#1 to #5) (**Fig 4B**). Notably, we observed a marked decrease in cMYC expression in the cells with stably knocked down FXR1 (**Fig 4B**). From these clones, we selected clone #1 for further functional assays. As expected, and in agreement with our siRNA results in **Fig 2**, knocking down FXR1 also significantly decreased cell proliferation and colony formation of HeyA8 cells (**Fig 4C and 4D**).

To determine the consequences of loss of FXR1 expression *in vivo*, we injected the control cells or FXR1-knockdown HeyA8 cells into the ovary bursa of nude mice orthotopically and monitored tumor progression (n=10 mice/group) for up to 5 weeks by bioluminescent imaging, using an *in vivo* imaging system (IVIS) (**Fig 4E**). Our IVIS imaging showed that FXR1 knockdown reduced ovarian tumor burden by ∼70% at the last three time points (**Fig 4F**). In a complementary analysis at the endpoint, we found that silencing FXR1 had markedly reduced tumor growth at both the primary site of injection and metastatic abdominal sites (**Fig 4G to 4I**). We also monitored the effects of changes in tumor burden on survival until all the mice in the shFXR1 group (n=8 mice/group) died (**Fig 4J**). Importantly, this analysis found that the mice injected with shFXR1-HeyA8 cells had survived longer than the shCont mice (median survival of 105 days vs. 33 days, log rank test *p* <0.0001) (**Fig 4K**). Our immunoblot analysis of the tumor lysates also showed that the tumors with stably knocked down FXR1 expressed lower levels of cMYC and Cyclin D1 compared to the controls (**Fig 4L**). Conversely, those tumors expressed higher levels of p21 and p27 proteins (**Fig 4L**). Next, we performed immunohistochemistry on tumor tissues collected from the mice. This showed that depletion of FXR1 decreased levels of the pro-survival marker Ki67 and our identified FXR1 target cMYC oncoprotein. We also found that stably knocking down FXR1 increased levels of cleaved caspase-3, a pro-apoptotic marker, compared to the control tumors (**Fig 4M**). Taken together, our results demonstrate that silencing FXR1 expression inhibits the growth and metastasis of HeyA8 ovarian cancer cells *in vivo*.

### Inducible expression of FXR1 increases levels of pro-oncogenic proteins and promotes oncogenic characteristics in ovarian cancer cells

Next, we created a simultaneous, efficient, and controlled model for inducing FXR1 by using a tetracycline (tet)-inducible vector (**Fig 5A**) to introduce the FXR1 gene fused to GFP into HeyA8 ovarian cancer cells. Cells that stably expressed inducible GFP-FXR1 were selected by FACS sorting after tetracycline treatment (1µg/ml) (**Fig 5A**). As expected, induction of FXR1 promoted the proliferation and colony formation of ovarian cancer cells compared to their control groups (**Fig 5B and 5C**). Our data also demonstrated that FXR1 induction by tetracycline increased levels of cMYC, Cyclin D1, CDK2, CDK4, and CDK6 proteins (**Fig 5D**).

We determined the effect of FXR1 on ovarian cancer progression *in vivo* by orthotopically injecting luciferase-tagged HeyA8 cells that stably expressed tet-inducible FXR1 (**Fig 5A**) into the left ovary bursa of (Foxn1/Nu) nude mice (n=8/group) (**Fig 5E**). To induce FXR1, we treated the mice with doxycycline (Dox, which is the Dox-derivative of tetracycline) in their drinking water (2mg/ml) throughout the experiment. We then monitored tumor growth through bioluminescent imaging (**Fig 5E**). Induction of FXR1 by Dox promoted the growth of tumors from the ovarian cancer cells at the primary injection site and increased the rate of metastasis to the other ovary and other organ sites (**Fig 5F to 5J, Videos-S1 and S2**). The enhanced expression of FXR1 upon Dox treatment was confirmed by immunoblot analysis, using lysates prepared from the tumor tissues. We found that the Dox-treated mice expressed high levels of FXR1 *in vivo* (**Fig 5K**) and that their tumor tissues contained increased levels of cMYC and Cyclin D1 (**Fig 5K**). Those tumors also expressed high levels of the proliferative marker Ki67 and low levels of the apoptosis marker cleaved caspase-3 compared to the mice not treated with Dox (**Fig 5L**). Taken together, these results complement our *in vitro* and *in vivo* data that suggest FXR1 silencing (**Fig 2 and 4**). They also confirm that overexpression of FXR1 promotes tumor growth and metastasis *in vivo*.

### Binding of FXR1 onto cMYC mRNA improves stability and enhances cMYC translation

Next, we sought to decipher the molecular mechanism that enables FXR1 to regulate cMYC levels in ovarian cancer cells. First, we tested the binding of FXR1 to *cMYC* mRNA, using an RNA immunoprecipitation (RIP) assay. The FXR1-bound mRNAs were pulled down with a monoclonal antibody specific to FXR1. Then we performed qPCR, using primers specific to *cMYC* mRNA (**Fig 6A**). Our results showed a significant increase (∼2 fold) in the enrichment of *cMYC* mRNA compared to the input (**Fig 6A**). In contrast, we did not find any detectable signal when we used control IgG (**Fig 6A**). Second, we tested the effect of FXR1 on *cMYC* mRNA translation with an *in vitro* translation assay, using the rabbit reticulocyte lysate system. As shown in **Fig 6B**, the yield of cMYC protein increased more (∼60%) in the presence of purified FXR1 protein than in its absence (**Fig 6B**). Third, we determined how FXR1 modulates the level of cMYC protein and its turnover rate. For this experiment, we transfected HeyA8 ovarian cancer cells with either control siRNAs or siRNAs specific to FXR1. The transfected cells were then treated with CHX for the indicated times, and cMYC protein levels were monitored. As shown in **Fig 6C**, inhibition of FXR1 resulted in rapid degradation of cMYC protein, with a ∼50% reduction in half-life. In a complementary approach, we performed a CHX-chase experiment while inducing FXR1 expression in HeyA8 cells. As expected, overexpression of FXR1 in HeyA8 cells improved the stability of cMYC protein compared to the control (∼68.5 min vs. 48.2 min) (**Fig S4A**). These results together demonstrate that FXR1 binds to *cMYC* mRNA and that its binding enhances cMYC translation.

**Fig 6:**
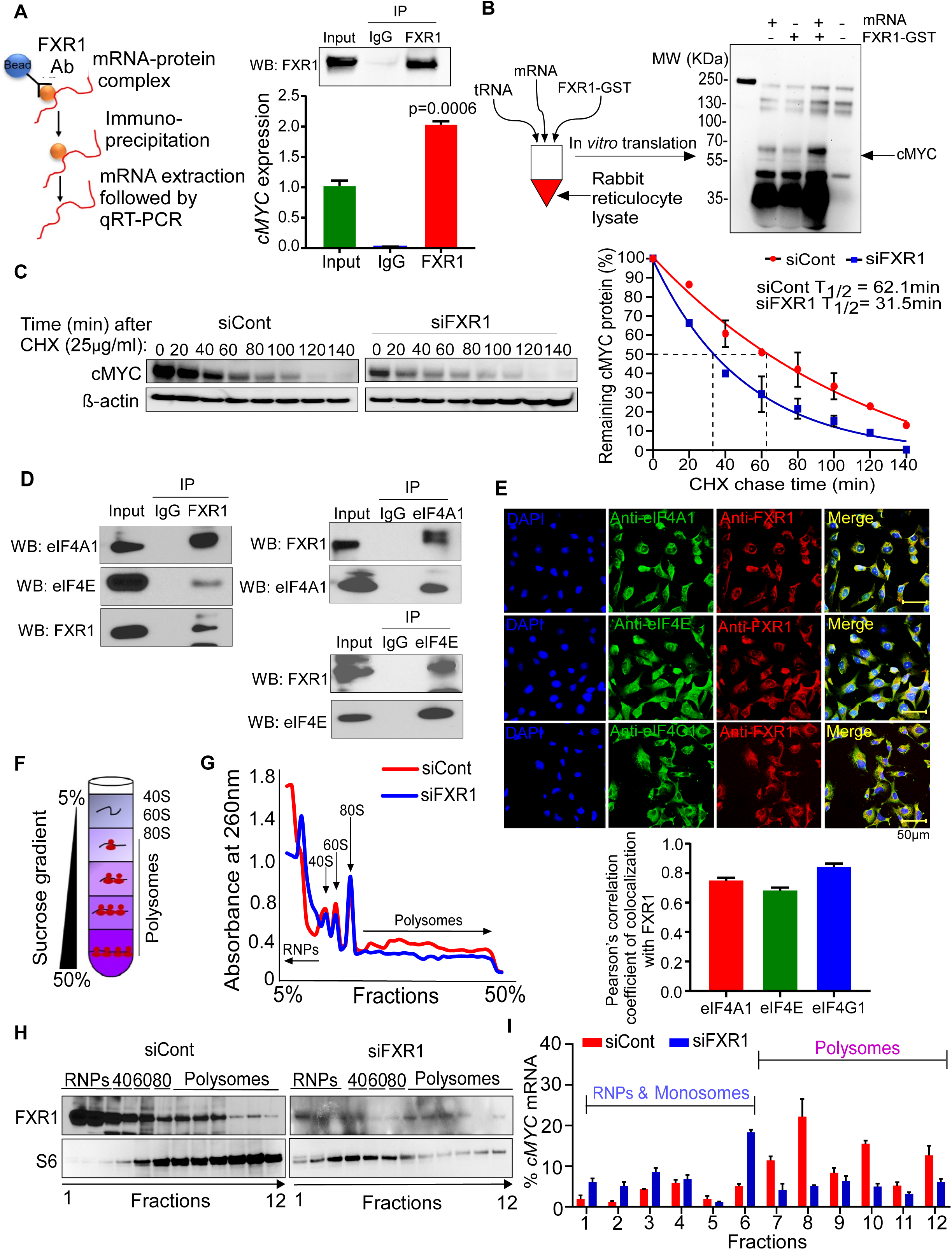
FXR1 stabilize cMYC post-transcriptionally and interacts with eukaryotic initiation factors (eIFs) for translation. **(A)** A schema presents how the RNA immunoprecipitation (RNA-IP) assay was performed using monoclonal antibodies specific to FXR1 (left). Total RNA was eluted from the RNA-IP and qPCR was performed for the enrichment of *cMYC* mRNA. Analysis was done by quantitating *cMYC* mRNA in the complex immunoprecipitated compared to control IgG (Bottom). Representative Western blot of FXR1 in the corresponding samples (Top). Error bars indicate SEM. ***p<0.001 compared to input by Student’s t-test. **(B)** *In vitro* translation assay was performed using the total RNA isolated from HeyA8 cells were incubated with FXR1-GST human recombinant protein for 1.5h, then immunoblotted using cMYC antibody. **(C)** Immunoblot was performed using the lysates prepared from the HeyA8 cells were transfected with siCont or siFXR1 for 48h, followed by the treatment with cycloheximide (25µg/ml) for indicated time points (left). Densitometric quantification measurements of cMYC protein bands from the blots were quantitated using image-J and analyzed for phase decay quantification (right). **(D)** HeyA8 cell lysates were immunoprecipitated using antibody specific to FXR1. The immunocomplexes were then eluted and immunoblotted using indicated antibodies. Normal immunoglobin G (IgG) was used as the negative control and 5% whole lysates (Input) was used as the positive control. **(E)** HeyA8 cell lysates were immunoprecipitated using antibodies specific to eIF4A1 or eIF4E. The immunocomplexes were then eluted and immunoblotted using indicated antibodies. Normal immunoglobin G (IgG) was used as the negative control and 5% whole lysates (Input) was used as the positive control. **(F)** HeyA8 cells were fixed with PFA, permeabilized and immune-stained with anti-FXR1, anti-eIF4s antibodies (eIF4A1, eIF4E, eIF4G1) and nuclear stain DAPI (top). Pearson correlation coefficient was assessed by calculating the co-localization signaling intensity using image-J Coloc-2 pixel intensity spatial correlation analysis suite. Error bars indicate SEM. **(G)** Schema represents how we performed the polysome profiling using sucrose gradient (5-50%) columns to collect fractions of RNAs bound to the ribosome units as indicated. **(H)** Plot of the absorbance profile of fractions obtained through sucrose gradients to isolate polysomes from HeyA8 cells transfected with control siRNAs or siFXR1 for 48h. Peaks and curves indicate the binding of RNA to the marked units of ribosome or polysome. **(I)** Western blot analysis of the protein fractions isolated from **H** using indicated antibodies. **(J)** qPCR shows the enrichment of *cMYC* mRNA in the isolated fractions bound with free RNPs, monosomes and polysomes. Data was normalized with the control group. Error bars indicate SEM.

We then sought to determine how FXR1 enhances the translation of *cMYC* mRNA into protein. Translation initiation is the rate-limiting step of protein production, which begins with the loading of the 43S pre-initiation complex (43S PIC, 40S ribosomal subunit associated with several initiation factors including a eIF2-GTP-Met-tRNA^Met^ ternary complex) onto the 5′mRNA cap bound to an eIF4F complex (consisting of the cap-binding proteins eIF4E/eIF4G, the helicase eIF4A, and the poly(A)-binding protein) [27]. Therefore, we determined whether FXR1 could associate with any components of the eIF4F complex. After preparing whole-cell extracts of the ovarian cancer cells and immunoprecipitating the extracts with monoclonal antibodies to FXR1, we found a strong interaction of FXR1 with both eIF4A1 and eIF4E (**Fig 6D**). When we used a reverse approach by immunoprecipitating eIF4A1 and eIF4E with target-specific monoclonal antibodies, we found that both eIF4A1 and eIF4E interact with FXR1 (**Fig 6E**). Consistent with these observations, our immunofluorescence assay showed co-localization of FXR1 with eIF4A1, eIF4E, or eIF4G1 in ovarian cancer cells (**Fig 6F)**. Collectively, these data suggest a strong association of FXR1 with proteins in the eIF4F complex. In conjunction, we also found that the levels of FXR1 protein correlated directly with eIF4F family proteins in ovarian cancer patients (**Fig S4B**). Again, this finding supports the notion that FXR1 could be important regulator of translation initiation.

In line with our results, a recent study reported that eukaryotic translation initiation factors (eIFs) interact with FXR family proteins [28]. Given that FXR1 is strongly associated with eIF4F proteins, we posited that FXR1 promotes the assembly of the initiation complex on the mRNA cap to promote protein synthesis. To test this idea, we knocked down FXR1 in HeyA8 cells and prepared cytoplasmic fractions, which were further fractionated on 5%–50% linear sucrose gradient columns. By recording absorbance of the fractions at 254 nm to profile polysome content, we obtained four peaks representing 40S and 60S ribosomal subunits, 80S monosomes and polysomes (left to right, **Fig 6G**). Notable, this assay showed a lower polysomal mRNA peaks for the FXR1-deficient cells. In contrast, monosomal peaks (80S) increased, which indicating inefficient translation or translational repression due to loss of FXR1 expression (**Fig 6H**). Therefore, we subsequently loaded the fractions we had collected from the gradients onto SDS-PAGE for Western blot analysis. In line with our previous results, knocking down FXR1 caused a striking shift of ribosomal protein RPS6 (S6) from polysomes to monosomes, while actively translating polysomes dominated in control cells (**Fig 6I**). As expected, all fractions contained FXR1 protein except the samples transfected with FXR1 siRNA (**Fig 6I**).

To quantify levels of *cMYC* mRNA bound to polysomes, we performed qPCR, using mRNA purified from the ribosomal fractions. Depletion of FXR1 reduced the amount of polysome-bound *cMYC* mRNA, confirming that FXR1 binds to *cMYC* mRNA (**Fig 6J**). Taken together, our results demonstrate that binding of FXR1 onto *cMYC* mRNA enhances translation by promoting the assembly of the initiation complex onto the mRNA cap.

### FXR1 directly interacts with AU-rich elements in the 3′UTR of cMYC for protein translation

Many RNA-binding proteins, such as IGF2BP3 [29], Mushashi [30], RBM3 [31], and FXR1[13] are known to bind to AU-rich elements (ARE) within the 3′ UTR of mRNA and to regulate translation in various cancer cells. Therefore, we sought to determine if FXR1 could bind to AREs within the 3′ UTR of *cMYC* mRNA to promote post-transcriptional changes. For this, we synthesized biotinylated RNA probes containing all six AREs spanning the entire *cMYC 3’UTR* (ARE1, ARE2, ARE3, ARE4, ARE5, and ARE6), and then performed RNA electrophoretic mobility shift assays (REMSA) (**Figs 7A and 7B, Table S5**). For negative controls, we used two random probes that lack ARE sequences, R1 and R2 (**Table S5**). The probes encompassing ARE1, ARE2, ARE4, and ARE5 showed strong binding with FXR1 protein but not with GST protein. In contrast, the other two probe targets exhibited weak (ARE3) or poor (ARE6) affinity with FXR1. Confirming the specificity of this interaction, incubation with excess amounts of cold probes decreased the binding of FXR1 with ARE (**Fig 7C**). Notably, no interactions were observed between the nonspecific R1 and R2 probes with FXR1 protein (**Fig S5A)**.

**Fig 7:**
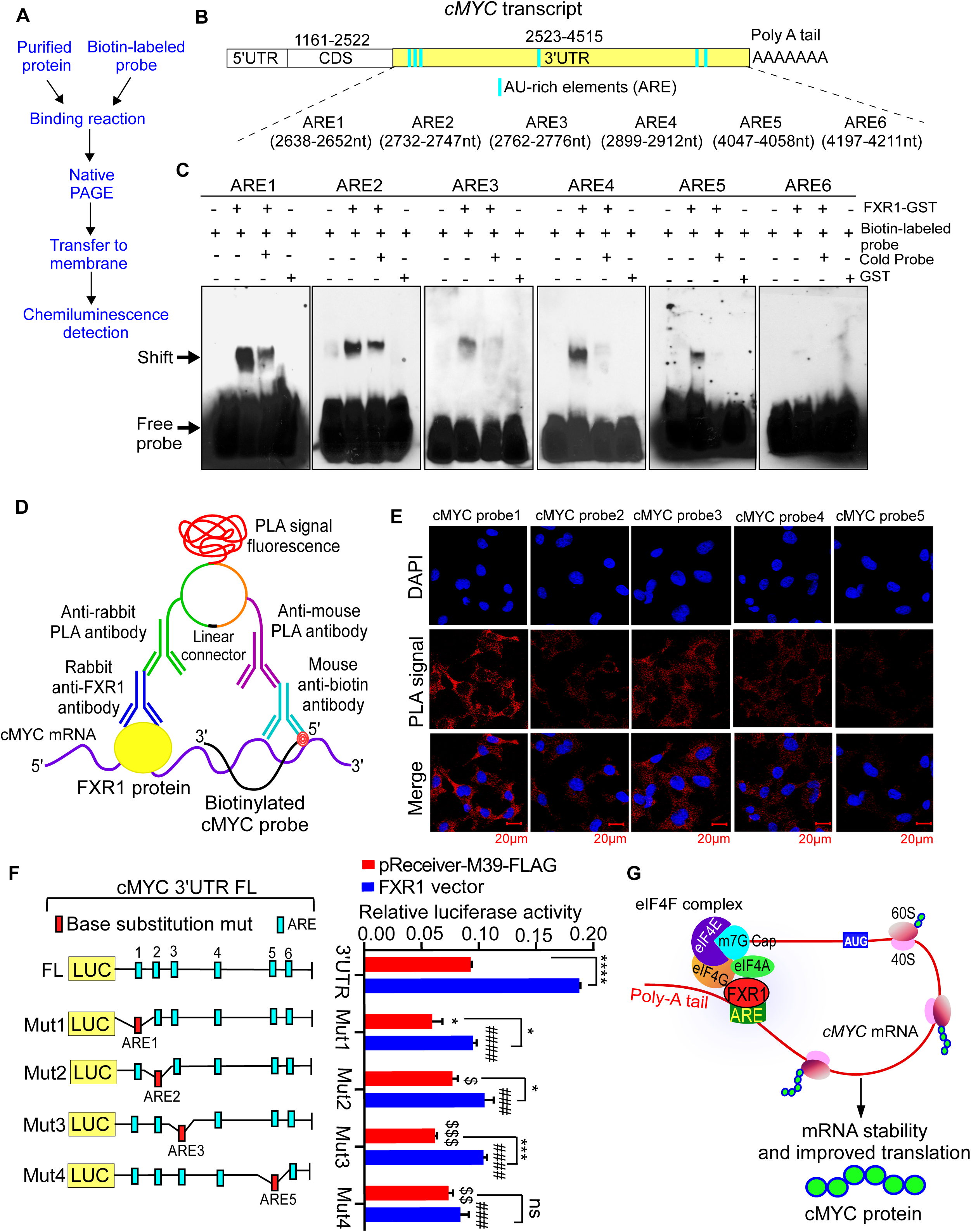
FXR1 binds to the AU-rich elements (ARE) in cMYC 3’UTR for translation. **(A)** Workflow for RNA electromobility shift assay (REMSA) shows FXR1 binding mediated mobility shift of *cMYC* mRNA. **(B)** Schema shows the number (ARE1 to ARE6) and position of ARE in the whole 3’UTR of *cMYC* transcript. **(C)** Biotinylated RNA probes containing 15-mer of human cMYC AREs were incubated with GST or FXR1-GST human recombinant protein for 30min. For competition assay, biotinylated RNA probes were mixed along with or without 200-fold of unlabeled cMYC probes. Reaction mixtures were then resolved on 6% gel and blotted to nylon membrane and developed by chemiluminescence assay. Arrow marks indicate the position of free probe and shift in the mobility due to FXR1 binding. **(D)** Schema of RNA-Proximity ligation assay (PLA) demonstrate the proximal localization of FXR1 protein with cMYC *mRNA*. **(E)** Representative confocal microscopy images (x63 magnification) of HeyA8 cells were fixed, permeabilized, and incubated with anti-FXR1 antibody and oligonucleotide probes anti-sense to 3’UTR of cMYC (1 to 5). PLA was then performed to detect the interaction of FXR1 protein and the *cMYC* mRNA (red); nuclei were stained with DAPI (blue). **(F)** Map of full length or mutated cMYC 3’UTR was prepared and cloned in the downstream of luciferase reporter in pEZX-MT06 vector, where red marks indicated base substitution mutations in the ARE sequence. **(G)** Each of the cMYC 3’UTR constructs in (F) were transfected in the OVCAR3 cells, which stably express FXR1 or control vector; then the luciferase reporter activity was measured 24h after transfection in the cell lysates using a plate reader. Results are shown as mean of three technical replicates ± SEM. ns: non-significant, *p < 0.05, ***p<0.001, ****p<0.0001 compared with empty control vector, ^###^p < 0.001, ^####^p<0.0001 compared to cMYC FL 3’UTR FXR1 vector, ^$^p<0.05, ^$$^p<0.01, ^$$$^p<0.001 compared to empty control vector by Student’s t-test. **(H)** Proposed model demonstrates that how FXR1 binding on 3’UTR of cMYC facilitate the translation by recruiting the eiF4F factors to translation initiation site.

To confirm that FXR1 binds with AREs in the 3’UTR of *cMYC* mRNA, we performed a RNA-proximity ligation assay (RNA-PLA), which detected close proximity between FXR1 protein and the ARE probes we identified in our REMSA (**Fig 7D)**. In this assay, we used five biotin-labeled PLA probes (probe1 to probe 5) that are 7 bases away from each ARE (**Fig S5B, Tables S5 and S6**). Using fluorescent signals, we detected proximal binding of FXR1 to the probe close to the ARE (**Fig 7D)**. As expected and complementary to our REMSA results, the *PLA Probe-1* (close to ARE1 and ARE2) and the *PLA Probe-3* (close to ARE4) provided the strongest fluorescent signals of proximity. *PLA Probes 2 and 4* (respectively close to ARE3 and ARE5) yielded moderate signals, while *PLA Probe 5* (close to ARE6) gave only weak signals (**Fig 7E and S5C for quantification**). As expected, the cells with FXR1 knockdown gave a poor signal compared to the cells transfected with control siRNA, even when we targeted their AREs using *PLA Probe-1,* which had given the highest yield for proximal ligation (**Fig S5D**). Taken together, our results demonstrate that FXR1 binds to target-specific sequences within the 3’UTR of *cMYC* transcripts.

To further confirm that FXR1 binds to the 3′ UTR of *cMYC* mRNA in a sequence-specific manner, we performed a luciferase reporter assay, using a construct of the wild-type 3′UTR of cMYC (FL-cMYC-3′UTR-Luc) containing all the ARE sequences cloned downstream of a dual (firefly and renilla) luciferase (Luc). We also used 3′ UTR mutants (Mut1 to Mut4 indicates a point mutation in each site, as indicated) (**Fig 7F, Table S7**). In this assay, we transiently transfected WT FL 3′ UTRs of the *cMYC* mRNA and four mutants (Mut1 to Mut4) into OVCAR3 cells that exogenously overexpressed FXR1. We then quantified cMYC 3’UTR reporter signals (**Fig 7F and 7G, Table S7**). As expected, overexpression of FXR1 along with FL-cMYC-3′ UTRs increased luciferase reporter activity as compared to the empty control vector, whereas the individual mutations in the ARE sequences in the 3’UTR reduced luciferase reporter activity compared to the FL-cMYC-3’UTR (**Fig 7G**). Moreover, there was lower luciferase reporter activity in the mutants that were transiently transfected with control vector (Mut1 to Mut4) than in those that overexpressed FXR1 (Mut1 to Mut4) (**Fig 7G**). These results demonstrate that FXR1 binds to the FL-cMYC-3′ UTR of *cMYC* transcripts, increasing protein translation. In light of these results, we hypothesize that binding of FXR1 onto the ARE in the 3’UTR of cMYC not only stabilizes the mRNA but also facilitates the recruitment and assembly of eukaryotic initiation factors (eIFs) to the translation initiation site. These events enhance the translational efficiency of *cMYC* mRNA, boosting the oncogenic potential of ovarian cancer cells, including growth and metastasis **(Fig 7H)**.

## DISCUSSION

RBPs are known for post-transcriptional actions that regulate the expression of many genes, including proto-oncogenes, apoptosis regulators, cell-cycle associated genes, and pro-inflammatory cytokines. These actions are regulated mainly by changing the decay kinetics of target mRNAs or altering translation to promote pathological conditions including cancer [32, 33].

Ovarian cancer is driven primarily by copy number variation or mutations in p53, BRCA1, or BRCA2 [1]. In this study, we report that a large subset of ovarian cancer patients expresses high levels of FXR1 due to CNV. Such patients suffered poor clinical outcomes compared with patients with lower FXR1 levels, suggesting that FXR1 is an important oncoprotein with a critical role in the pathophysiology of ovarian cancer. Using both *in vitro* and orthotopic mouse models, we confirmed that FXR1 promoted oncogenic effects in ovarian cancer, including hyperproliferation of tumor cells and tumor cell invasion and metastasis. Our mechanistic and correlative clinical data indicate that overexpressed FXR1 behaves as an oncogene by upregulating cMYC in ovarian cancer. We further found that FXR1-mediated cMYC upregulation promoted the levels of Cyclin E1, Cyclin D1, and CDKs along with extensive proliferation of ovarian cancer.

The cMYC oncogene is one of the commonly upregulated oncogenes in human cancers, being dysregulated in more than 40% of cases, and its high expression frequently associates with poor prognosis and unfavorable patient survival [34, 35]. On this context, our study established an important mechanism that promotes the protein synthesis of cMYC protein. Our pathway and gene set enrichment analyses along with biochemical approaches showed that FXR1 dysregulates a variety of pathways that are critical for tumor progression. We also found an association between FXR1 and cMYC in ovarian cancer patient samples. Our data proved that cell cycle changes promotes the proliferation, as well as metastatic effects of FXR1 operate through the gain in cMYC protein levels.

Mechanistically, we provided evidence that FXR1 binds onto *cMYC* mRNA. Specifically, we found that FXR1 binds to specific sequences of ARE (AUUUA) within the cMYC 3′ UTR. We further proved that binding of FXR1 onto *cMYC* mRNA improved the rate of translation. In complement, FXR1 improved the enrichment of polysomes on *cMYC* mRNA as a critical mechanism for enhanced cMYC translation. Using both immunoprecipitation and co-localization assays, we verified that the interaction between FXR1 and eIF4F proteins is direct, which again supports the notion that FXR1 boosts the synthesis of cMYC protein. The interaction between eIF4F proteins and the 7-methylguanosine “cap” (m^7^G) located at the 5′ end of all mRNAs is critical to directly recruit the 40S ribosomal subunit to mRNAs through a set of protein–protein interactions and to unwind RNA secondary structures in the 5′ untranslated region (5′ UTR) of mRNAs [36]. Our data suggest that the FXR1–eIF4F loop promotes translation, presumably by recycling ribosomes as proposed for eIF4G–PABPC1-mediated mRNA looping, which was established previously [37]. Compatible with published data, our data also suggest that looping between the stop codon and the 5′ UTR end might be a more efficient way to recycle ribosomes rather than via the 3′ UTR end, especially for mRNAs with long 3′ UTRs, as ribosomes would dissociate from the mRNP once ribosomes were released at the stop codon.

A recent study demonstrated that FXR1 promotes prostate cancer progression by directly associating with *FBXO4* 3’UTR and thereby upregulating mRNA stability [11]. Another report showed that FXR1 promotes the malignant biological behavior of glioma cells by stabilizing MIR17HG [12]. However, we did not find any correlation between FXR1 and FBXO4 or MIR17HG in our clinical samples of ovarian cancer. Therefore, FXR1 CNV may have selective, multifunctional effects that depend on the repertoire of target oncogene transcripts expressed in given tumors. This needs to be investigated, as does the complexity of FXR1’s actions and its precise role in cMYC regulation in different cancer types. Because FXR1 is highly amplified and expressed in many types of cancers, it may have widespread oncogenic roles. Secondly, direct targeting of cMYC has been a challenge for decades owing to its “undruggable” protein structure. From a clinical perspective, we believe that FXR1 may represent a new target that holds great potential for suppressing cMYC expression in malignancies that exhibit FXR1 CNV. Such studies on inhibiting FXR1 for therapeutic purposes are currently undergoing in our laboratory.

## Conclusion

Our study uncovered a novel mechanism that enables FXR1 CNV to upregulate the cMYC oncogene in ovarian cancer. Our *in vivo* data using an orthotopic ovarian cancer model demonstrated that inhibiting the actions of FXR1 in cancer cells that exhibit FXR1 CNV could be a viable approach to developing therapeutic strategies in ovarian cancer. Moreover, significant upregulation of FXR1 expression correlates with poor prognosis of patients who have the disease. Thus, FXR1 alone or in combination with its downstream effectors could be an important biomarker in ovarian cancer.

## MATERIALS AND METHODS

### Patient Samples

After informed consent, normal ovarian epithelial tissues and serous epithelial ovarian cancer tissues were collected according to an Institutional Review Board approved protocol at Medical College of Wisconsin. Normal ovarian tissues were from the tumor-free ovary of patients with unilateral ovarian cancer or patients with other gynecological cancers not involving the ovary.

### Cell culture

OVCAR3, SKOV3, OVCAR5, OVCAR8 and A2780 were purchased from the National Cancer Institute (NCI) cell line repository. HeyA8 cells were received from the Characterized Cell Line core at MD Anderson Cancer Center, Houston, TX, USA. PEO1 cells were received from Daniela E Matei, Northwestern University, Chicago, Illinois, USA. Immortalized fallopian tube epithelial cells FTE188 was received from Jinsong Liu and MCAS was received from Gordon Mills at MD Anderson Cancer Center, Houston, Texas, USA. All cancer cell lines were cultured in DMEM medium (Sigma-Aldrich, Saint Louis, MO) supplemented with 10% fetal bovine serum (FBS, Atlanta Biologicals, GA, USA), penicillin (100 U/ml) and streptomycin (0.1 mg/ml) and Sodium pyruvate (Sigma-Aldrich). FTE188 was maintained in cell culture medium consisting of 1:1 Medium 199 and MCDB105 medium (Sigma-Aldrich) with 10% FBS and 10ng/ml EGF (Sigma-Aldrich). Cells were routinely tested and deemed free of Plasmotest^TM^ Mycoplasma Detection Kit (InvivoGen, San Diego, CA). Authenticity of the cell lines used were confirmed by STR characterization at IDEXX Bioanalytics Services (Columbia, MO).

### Tumor models

Female athymic nude mice (CrTac: NCr-Foxn1nu, Taconic Laboratories, RRID: IMSR_TAC:ncrnu), approximately 4 to 6 weeks old were maintained under specific pathogen-free conditions in accordance with guidelines and therapeutic interventions approved by the Institutional Animal Care and Use Committee (IACUC) at the Medical College of Wisconsin. For tumorgenicity study, HeyA8 cells with stable knockdown of FXR1 (shFXR1) and control cells (shCont) were trypsinized, washed and resuspended in Hanks’ balanced salt solution (Gibco, Carlsbad, CA), then a total number of 25,000 cells/mice in culture medium were injected orthotopically into the ovarian bursa of the anesthetized female nude mice (n=10) through a 1.5-cm intraperitoneal incision as described before[38]. For survival study, shCont and shFXR1 cells were injected intraperitoneally in mice (n=8). Similarly, HeyA8 cells with stable Tet-induced FXR1 overexpressed (Tet (+)) and control (Tet (-)) cells were processed and injected into mice ovary orthotopically (n=8) followed by Doxycycline (2mg/ml) and 10% (w/v) sucrose (Sigma-Aldrich) in drinking water after 1 weeks of cells inoculation. The solution was protected was freshly prepared every second day. Tumor tissue was prepared as snap frozen for protein isolation or fixed in 10% formalin for immunohistochemistry.

### Tissue Micro-Array (TMA) and immunohistochemistry (IHC)

FXR1 protein levels in human ovarian cancer tissues were analyzed using two TMAs (Cat# OV1005bt and Cat# OVC961, US Biomax Inc., Rockville, MD) and cMYC protein levels in one TMA (Cat# OVC961). For this purpose, the slides were dewaxed in xylene, and rehydrated through graded ethanol to distilled water. Antigen retrieval for the slide specimens were performed using IHC-Tek epitope retrieval solution and steamer set (IHC World, LLC.). The slides were then immersed in 3% H_2_O_2_ for 10 min to quench endogenous peroxidase followed by blocking with 10% goat serum for 1 h. Vectastain ABC-AP Kit (Vector Labs, Burlingame, CA) and Vector Red Alkaline Phosphatase Substrate Kit I (Vector Labs, Burlingame, CA) were used for tissue staining as per manufacture protocol. FXR1 primary antibody (Proteintech, Cat#13194-1-AP) was used at 1:200 dilution and cMYC primary antibody (Santacruz Biotechnology, Cat#sc-47694) at 1:100. Following, Vector red staining, the slides were counterstained with Harris modified hematoxylin (Thermo Fisher Scientific Inc., Rockford, IL), dehydrated with graded ethanol and xylene, and finally mounted with paramount. TMAs slides was digitally scanned using Pannoramic 250 FLASH III scanner (3D HISTECH ltd. Version 2.0) and, using the Case Viewer software (3D HISTECH ltd. Version 2.0) was used to view and analyze images.

### Western blotting

For preparing cell lysates, the cells were washed twice with ice-cold PBS and lysed on ice in 1x RIPA lysis buffer containing freshly added protease inhibitor cocktails (Thermo Fisher Scientific Inc., Rockford, IL, USA) and 1 mM PMSF. For preparing the tissue lysates, the tumor tissues were homogenized in 1x RIPA lysis buffer over ice. After 30 min of incubation, the lysates were collected by centrifugation at 4°C for 10 min at 10,000 rpm. The amount of total protein was determined using a BCA protein assay kit (Pierce, Rockford, IL, USA). An equal amount of total protein (30μg) were resolved on precast 4-12% SDS-PAGE gels (Biorad, Hercules, CA, USA), transferred onto nitrocellulose membranes, and incubated with desired primary antibodies, followed washing and incubation with HRP conjugated secondary antibodies and detecting of protein bands using chemiluminescence kit (Pierce, Rockford, IL, USA).

### siRNA transfection and cell viability assay

Predesigned siRNAs for human FXR1 were purchased from Thermo Fisher Scientific Inc., Waltham, MA and negative siRNAs universal control (siCont) from Sigma-Aldrich. Transfections were performed using the Lipofectamine RNAiMAX transfection reagent (Thermo Fisher Scientific Inc., Waltham, MA). At 48 h post-transfection, cells were harvested for further analysis. siRNA sequences are listed below:

FXR1 siRNA (siFXR1-1):
Sense: GGAAUGACUGAAUCUGAUAtt
Antisense: UAUCAGAUUCAGUCAUUCCat FXR1 siRNA (siFXR1-2):
Sense: CGAGCUGAGUGAUUGGUCAtt
Antisense: UGACCAAUCACUCAGCUCGtc

### Cell proliferation and colony formation assays

Cell viability was measured with the Cell Counting Kit-8 (CCK-8) (Dojindo, Shanghai, China) according to the manufacturer’s instructions. Cells were plated at a density of 1×10^3^ cells per well in 96-well plates and incubated at 37°C. Proliferation rates were determined at 0, 24, 48, 72 h post-transfection, and quantification was performed on a microtiter plate reader (Tecan, Mannedorf, Switzerland) at 450nm.

For colony formation assay, transfected cells were plated in six-well plates at a density of 1000 cells per well. After 10 days, cells were rinsed with PBS, fixed in 5% glutaraldehyde for 10min and then stained with 0.5% crystal violet (Sigma Aldrich, MO, USA) for 20min. Plates were washed with water and dried before imaging.

### Cell invasion assay

Ovarian cancer cells were treated with siCont and siFXR1s for 12h followed by re-plating of treated cells (2 x 10^5^ cells) to the matrigel (Corning NY, USA) coated inserts in the presence of cell cycle inhibitors mitomycin C (5 μg/ml). Cells were incubated at 37 °C for 24h for invasion assays. Cells that did not invade through the pores were removed using a cotton swab and inserts were washed and stained with 0.5% crystal violet for imaging. Alternatively, stained membranes were dissolved in 10% acetic acid, and quantified in microplate reader (Tecan, Mannedorf, Switzerland) at 560 nm.

### Quantitative real time-PCR (qRT-PCR)

Total RNA was isolated from the cells using RNeasy Mini Kit (Qiagen, Valencia, CA, USA) and first strand cDNA was transcribed with Oligo(dT) primers, dNTPs and M-MLV reverse transcriptase (Promega, Fitchburg, WI, USA). qRT-PCR was performed using CFX Connect Real-Time PCR systems (Biorad, Hercules, CA, USA) and SYBR Premix Ex Taq II (Biorad, Hercules, CA, USA) with first strand cDNA, forward and reverse primers. The following primers were used: *FXR1*, Forward: 5’-GAGAGAAGATTTAATGGGCCTGG-3’ and Reverse: 5’-GCTCAATGGCGGTAACTCCA-3; *GAPDH*, Forward: 5’-CTCACCGGATGCACCAATGTT-3’ and Reverse: 5’-CGCGTTGCTCACAATGTTCAT-3’. The PCR program used was as follows, initial denaturation step (95°C for 30 s) followed by DNA amplification (95°C for 3 s followed by 60°C for 30 s) for 40 cycle. Melt curve analysis was performed to ensure the specificity of target amplicon. Relative mRNAs were analyzed using *GAPDH* as endogenous control and ΔΔCT algorithm. PCR array for human cell cycle (Cat#PAHS-020Z) and oncogenes and tumor suppressor genes (Cat#PAHS-502Z) was purchased from Qiagen (Valencia, CA, USA), and was used according to the manufacturer’s instructions. Data for PCR array were analyzed using SABiosciences RT^2^ Profiler PCR Data Analysis software at http:// pcrdataanalysis.sabiosciences.com/pcr/arrayanalysis.php and were considered significant at > 1.5 fold change and p<0.05. Five housekeeping genes, *B2M*, *HPRT1*, *RPLP0*, *GAPDH*, and *ACTB*, were used for normalizing data and fold change was calculated relative to the siCont treated HeyA8 cells.

### Cell cycle analysis

siFXR1s transfected ovarian cancer (OVCAR5 and HeyA8) cells were seeded at a density of 5 x 10^5^ in 6-well plates. When cells reached 70–80% confluence, they were washed with PBS, trypsinized, collected and fixed with 70% ethanol overnight. Next day, cells were treated with 1 mg/ml RNase A (Sigma-Aldrich, Saint Louis, MO) at 37°C for 30 min and then resuspended in 0.5 ml of PBS and stained with 50 μg/ml propidium iodide (PI) (Sigma-Aldrich, Saint Louis, MO). The cells were analyzed using a FACScan flow cytometer (Becton–Dickinson, Mansfield, MA) and ModFit LT software (Verity Software, Topsham, ME).

### Live/dead cell assay

Cellular death was measured with Live-or-Dye™ 488/515 Fixable Staining Kit (Biotium, USA) as per as manufacturer’s protocol. In brief, HeyA8 and OVCAR5 cells were reverse transfected with siRNAs (siCont and siFXR1). After 48h, the cells were washed with PBS and dye was added to the cells followed by incubation for 30min. Cells were fixed in 4% PFA, permeabilized with 0.5% Triton X-100 and stained with DAPI. Fluorescence images were acquired using a confocal laser scanning microscope (LSM 510; Zeiss, Oberkochen, Germany).

### Annexin V/PI staining for apoptosis

Cellular apoptosis was measured with FITC Annexin V Apoptosis Detection Kit I (BD Pharmingen, San Diego, CA, USA) as per as manufacturer’s protocol. In brief, HeyA8 and OVCAR5 cells were reverse transfected with siRNAs (siCont and siFXR1). After 48h, the cells were trypsinized, washed with PBS and resuspended in Annexin V binding buffer at a concentration of 10^6^ cells/ml. Annexin V– FITC (5μl) was added, vortex-mixed gently and incubated for 15 min at 4 °C in the dark. Cells were stained with 5 μl of PI for another 5 min at 4 °C in the dark. Stained cells were acquired on a FACS Calibur flow cytometer (Becton–Dickinson, Mansfield, MA) and data were analyzed with Flowjo software version 10.6.1 (TreeStar, Ashland, OR, USA).

### Caspase3/7 activity assay

Following treatments with siFXR1s for 48h, cells were subjected to Caspase 3/7 activity measurement with Caspase-Glo assay kit (Promega, Madison USA). Briefly, the plates containing cells were removed from the incubator and allowed to equilibrate to room temperature for 30 minutes. 100 μl of Caspase-Glo reagent was added to each well, the content of well was gently mixed with a plate shaker at 300–500 rpm for 30 seconds. The plate was then incubated at room temperature for 2 hours. The luminescence of each sample was measured in SpectraMax i3x Multi-Mode Microplate Reader (Molecular Devices, Japan Co., Ltd., Tokyo, Japan).

### Human apoptosis protein array

A human apoptosis array (Proteome Profiler™ Cat# ARY009; R&D Systems) was used to analyze apoptosis-related protein profiles according to manufacturer instructions. In brief, the total HeyA8 cell lysates after siFXR1-2 treatment (48h) were first incubated with the array membrane overnight at 4°C, followed by incubation with a biotinylated detection antibody cocktail at room temperature for 1 h. The membranes were then exposed and quantified by Image J software (National Institutes of Health, Bethesda, USA).

### Lentiviral FXR1 knockdown

For viral creation, HEK293T cells were transfected with packaging vectors pLP1, pLP2 and VSVG plasmids including control empty vector pLKO.1 (Cat#SHC001V) and two different FXR1 targeting short hairpin RNA (shRNA) (TRC number 1: TRCN0000160812, Clone ID:NM_005087.1-130s1c1; 2: TRCN0000160901, Clone ID: NM_005087.1-579s1c1) purchased from Sigma-Aldrich (Saint Louis, MO). Competent lentiviruses were collected 48 h after transfection. HeyA8 cells were passaged to 40% confluence, the next day viral media were added to cells with 8 μg/ml of polybrene. Efficacy of individual FXR1 shRNA construct was checked by Western blot analysis for FXR1 knockdown using FXR1 antibody (Cat#12295, Cell Signaling Technology). The most effective shRNA construct was used for generating FXR1 knockdown stable cell line by selection with puromycin (8 μg/ml; for 2 weeks). The clones were picked and subjected to expansion culture under further selection. Western blot analysis was performed to identify the stable clone with most efficiently down-regulated FXR1 protein, which was used in further experiments.

### Transfections for forced and inducible FXR1 overexpression

To establish the stable overexpression of FXR1 in OVCAR3 cells, we transfected the cells with control vector or pReceiver-M39 vector expressing FXR1 (GeneCopoeia, Rockville, MD) using Lipofectamine 2000 (Invitrogen, Carlsbad, CA). Cells were selected 48h after transfection to ensure that the cells were stably incorporated with control sequences or FXR1 using puromycin (2.5 μg/ml) containing culture media for two weeks. Western blotting was performed to check the expression of FXR1 and other targets in stable cells followed by colony formation assay for 12 days.

In order to create the tetracycline inducible cell line, HeyA8 cells were plated in a 6-well dish and transfected with pTRE-Tight GFP FXR1-expressing plasmid constructed at Vector Core facility, Versity Blood Research Institute using lentivirus method as mentioned above. GFP positive cells were selected by FACS sorting after tetracycline (Takara Bio Inc., San Francisco, CA, USA) treatment (1µg/ml). Transduced cells were expanded and used for further experiments.

### RPPA assay and data processing

Reverse Phase Protein Array (RPPA) and data analysis was performed as previously described [26] and detailed at the MD Anderson Cancer Center RPPA core facility as below: https://www.mdanderson.org/research/research-resources/core-facilities/functional-proteomics-rppa-core.html.

Briefly, cells were washed in ice-cold PBS, and lysed in 30 μ L of RPPA lysis buffer [1% Triton X-100, 50 nmol/L HEPES (pH 7.4), 150 nmol/L NaCl, 1.5 nmol/L MgCl_2_, 1 mmol/L EGTA, 100 nmol/L NaF, 10 nmol/L NaPPi, 10% glycerol, 1 nmol/L PMSF, 1 nmol/L Na_3_VO_4_, and protease inhibitor cocktail] for 30 minutes with frequent vortexing on ice, followed by centrifuging for 15 min at 14,000 rpm, and the supernatant were collected. Protein concentration was determined by Pierce BCA Protein Assay kit (Thermo Fisher Scientific, Waltham, MA) with a BSA standard curve according to manufacturer’s protocol. 30μL lysates were transferred into a 96-well PCR plate. To each sample well, 10μL of SDS/2-ME sample buffer (35% glycerol, 8% SDS, 0.25 mol/L Tris-HCl, pH 6.8; with 10% β-mercaptoethanol) was added and incubated for 5 minutes at 95°C and then centrifuged for 1 minute at 2,000 rpm.

Samples were diluted serially and transferred into 384-well plates and heated at 95°C for 10 min. Approximately, 1 nL of protein lysate was then printed onto nitrocellulose-coated glass slides (FAST Slides, Schleicher & Schuell BioScience, Inc., Keene, NH) with an automated robotic GeneTac arrayer (Genomic Solutions, Inc., Ann Arbor, MI) per array by pin touch. Each spot on the array slide represents a certain dilution of the lysate of a sample. Following slide printing, the array slides were blocked for endogenous peroxidase prior to the addition of the primary antibody, then treated with biotinylated secondary antibody (anti-mouse or anti-rabbit) was used as a starting point for signal amplification. Tyramide-bound horseradish peroxidase cleaves 3,3′-diaminobenzidine tetrachloride, giving a stable brown precipitate with excellent signal-to-noise ratio. Signal intensity was captured by scanning the slides with ImageQuant (Molecular Dynamics, Sunnyvale, CA) and quantified using the MicroVigene automated RPPA module unit (VigeneTech, Inc., North Billerica, MA). The intensity of each spot was calculated, and an intensity concentration curve was calculated with a slope and intercept using MicroVigene software.

### Protein decay analysis

For cMYC protein stability experiments, after treatment of HeyA8 cells with siCont and siFXR1-2 for 48 h, 25 μg/ml of CHX was added to inhibit protein synthesis and samples were collected at every 20 minutes for 140 min. Cells were then processed as previously described for Western blotting and quantified by densitometry.

### RNA immunoprecipitation (RIP) assay

RIP assay was performed using Magna RIP™ RNA-Binding Protein Immunoprecipitation Kit (Millipore Sigma, Burlington, MA) according to manufacturer’s instructions. In brief, following whole cell protein extraction from HeyA8 cells (2 × 10^7^), lysates were incubated with respective antibodies coupled to Dynabeads Protein A/G for overnight at 4°C. Following extensive washes, the immobilized immunoprecipitated complexes were incubated with proteinase K at 55°C for 30 min to digest the protein. Co-precipitated RNA and the Input (crude lysate) were eluted and purified with Trizol Reagent and analyzed by qPCR. The following primer was used for *cMYC*, Forward: 5’-AAACACAAACTTGAACAGCTAC-3’ and Reverse: 5’-ATTTGAGGCAGTTTACATTATGG-3.

### *In vitro* translation assay

*In vitro* translation assay for cMYC was performed by rabbit reticulocyte lysate system (Promega, USA) according to manufacturer’s instructions. In brief, total RNA (1µg/µl) from HeyA8 cells was subjected to *in vitro translation* by addition to 35 μl of rabbit reticulocyte lysate, methionine-free amino acid mixture, 40U of RNasin, transcend tRNA, and 4 μ g of purified human recombinant FXR1-GST protein (Novus Biologicals) for 1.5 h at 30°C. The reaction mixture was resolved by SDS-PAGE and cMYC translation was checked by Western blot analysis using cMYC antibody (Cat# 5605S, Cell Signaling Technology).

### SUnSET assay

HeyA8 cells were transfected with siFXR1-2 and after 48 hours incubated with 10µg/ml puromycin (Cat# A1113803, Thermo Fisher Scientific) in time dependent manner. Cells were washed with PBS to remove residual puromycin and collected by scraping. Cells were lysed using 1x RIPA buffer lysates and run on an SDS-PAGE gel, transferred to a nitrocellulose membrane for Western blotting with anti-puromycin antibody (Millipore, Cat#MABE343). Puromycin staining in lanes was measured using chemiluminescence kit (Pierce, Rockford, IL).

For flow cytometry, HeyA8 cells were transfected with siCont or siFXR1-2 for 48h and treated with CHX (50µg/ml) for 1h followed by with incubation with 10µg/ml puromycin for 10min. Cells were washed with PBS to remove residual puromycin followed by staining of cells using Alexa fluor-488 anti-puromycin antibody (Millipore, Cat#MABE343-AF488).

### Co-immunoprecipitation

For immunoprecipitation, Dynabeads® Co-immunoprecipitation Kit including Dynabeads® M-270 Epoxy beads were used (Thermo Fisher Scientific, MA, USA). 5mg of beads/IP were conjugated with respective antibodies (FXR1 Proteintech, Cat#13194-1-AP), eIF4A1 Sigma-Aldrich, Cat# SAB2700953, eIF4E, Sigma-Aldrich Cat# E5906, eIF4G, Santa Cruz Biotechnology Cat# 3344) overnight at 37°C with shaking. Antibody conjugated beads were added to cell lysates and allowed to incubate on a rotator for four hours at 4°C. Following incubation, beads were removed and washed with cold lysis buffer, boiled and analyzed by Western blot.

### Immunostaining

Immunostaining was performed as described previously [5, 38]. Briefly, HeyA8 cells were grown on eight-well chamber slides (ibidi USA, Madison, WI, USA) then fixed with 4% paraformaldehyde and permeabilized with 0.1% Triton X-100 in blocking solution (0.5% BSA in PBS) followed by blocking for 1 h with 0.5% BSA in PBS, and then stained overnight at 4°C with the indicated primary antibodies. After washing with PBS, cells were incubated with secondary antibodies, Alexa Fluor goat anti-mouse 488 (Cat#38731, Life Technologies, Carlsbad, CA) and Alexa Fluor 568 goat anti-rabbit (Cat#35646, Life Technologies, Carlsbad, CA) for 1 hour at room temperature. Glass slides were mounted using ProLong Gold Antifade Reagent (Life Technologies, Carlsbad, CA) containing DAPI. Images were acquired with a 40X objective using a confocal laser scanning microscope (LSM 510; Zeiss, Oberkochen, Germany) and analyzed using the Aim 4.2 software LSM510.

### Polysome fractionation by sucrose gradients

Polysome fractionation was performed following a previously published protocol [39]. In brief, HeyA8 cells were transfected with siFXR1-2 for 48hrs and total lysate was prepared by scraping in hypotonic buffer (5 mM Tris-HCl, pH 7.5, 2.5 mM MgCl_2_, 1.5 mM KCl and 1x protease inhibitor cocktail) supplemented with 100 μ g/ml cycloheximide (CHX), 1 mM DTT, and 100 units of RNAse inhibitor. Triton X-100 and sodium deoxycholate were added to a final concentration of 0.5% each, and samples were vortexed for 5 sec. Samples were centrifuged at 16,000 g for 7 min at 4°C. Supernatants (cytosolic cell extracts) were collected and absorbance at 254 nm was measured. Approximately 10-15 OD260s of lysate was layered over 5% – 50% cold sucrose gradients in buffer (200 mM HEPES (Ph 7.6), 50 mM MgCl_2_, 1 mM KCl, 100 μ g/mL CHX and 1x protease inhibitor, 100 units of RNAse inhibitor). Gradients were centrifuged at 39,000 rpm in a Beckman SW28 rotor for 2 h at 4°C. After centrifugation, 14 equal-sized fractions (0.75 mL/fraction) were collected and analyzed through UV detection. For Western blotting, fractions were precipitated with 95% alcohol and mixed with SDS sample buffer. For qPCR of cMYC, mRNA was isolated from the fractions by mixing with phenol:chloroform:isoamyl alcohol (Sigma-Aldrich) and then RNA was analyzed by qRT-PCR .

### Bioluminescence optical imaging

The IVIS Lumina II Bioluminescence and Fluorescence Imaging System (Caliper Life Sciences) was used for *in vivo* bioluminescent imaging. Mice were injected (i.p.) with 150 mg/kg body weight D-luciferin substrate (Gold Biotechnology, St. Louis, MO) and imaging was performed 10 min later (the peak time point). Images of the tumor were taken under the following settings: exposure time = 0.5 seconds, f/stop = 16, medium binning, field of view = 12.5 × 12.5 cm^2^. Living Image software was used to quantify the bioluminescent signals, reported as units of tissue radiance (photons/s/cm^2^/sr).

To further obtain the whole body three-dimensional (3D) images of mice to monitor tumor growth and metastasis, fluorescence Positron Emission Tomography-computed tomography (PET-CT) imaging (Perkin Elmer, Shelton, CT) was performed and images/videos were captured using Living Image software.

### RNA electrophoretic mobility shift assay (REMSA)

Biotin labelled and unlabeled RNA oligonucleotides probes corresponding to the human cMYC ARE (1-6) and two random probes (R1, R2) were synthesized as indicated in Table S5 by Sigma-Aldrich. REMSA was performed with a LightShift chemiluminescent RNA EMSA Kit (Thermo Fisher Scientific) following the manufacturer’s instruction. Briefly, purified human FXR1-GST protein (5 mg/ml) (Novus Biologicals, USA) and purified human GST protein (5 mg/ml) (Sigma-Aldrich) were incubated for 30min with biotinylated probes (100 pM) in REMSA binding buffer and glycerol. In competition binding assays, unlabeled RNA oligonucleotides (Sigma-Aldrich) were added in increasing amounts (200-fold) and incubated for another 10 min at room temperature. RNA/protein complexes were then electrophoreticed by 6% native polyacrylamide gel and transferred to nylon membrane (Thermo Fisher Scientific). RNA was cross-linked with a UV lamp at a distance of 0.5 cm from the membrane for 2 min. The membrane was blocked in blocking buffer for 15 min and replaced the blocking buffer with conjugate/blocking buffer. After washed with 1× wash buffer for 3 times, membrane was incubated in substrate equilibration buffer for 5 min. Then, the membrane was incubated in working solution and exposed.

### Proximity ligation assay

The DuoLink® In Situ Red Starter Kit Mouse/Rabbit (Sigma-Aldrich) was used to detect proximity between FXR1 protein and *cMYC* mRNA according to manufacturer protocol. Briefly, HeyA8 cells were seeded in eight-well chamber slides (ibidi USA, Madison, WI, USA) and cultured overnight. Slides were washed with cold 1×PBS and fixed in 4% paraformaldehyde for 30 min. Then slides were blocked with Duolink Blocking Solution in a pre-heated humidified chamber for 1hr at 4°C and followed by hybridizations with probes targeting cMYC RNAs (**Table S6**), respectively at 37°C. The primary antibodies to detect FXR1 (Proteintech, Cat#13194-1-AP) and biotin (Cat# 07-599, Rockland) was added to the slides and incubated overnight at 4°C. Then slides were washed with 1×Wash Buffer A and subsequently incubated with the PLA probes (1:5 diluted in antibody diluents) for 1h, then the Ligation-Ligase solution for 30 min, and the Amplification-Polymerase solution for 100 min in a pre-heated humidified chamber at 37°C. Before imaging, slides were washed with 1×Wash Buffer B and mounted with a cover slip using Duolink In Situ Mounting Medium with DAPI. Fluorescence images were acquired using a confocal laser scanning microscope (LSM 510; Zeiss, Oberkochen, Germany).

### Dual-luciferase reporter assay

The full-length (FL) cMYC 3’UTR (NCBI Reference Sequence: NM_001354870.1; 3’UTR region: 2523-4515) and four mutants (M1-M4) with base substitution in certain AREs cloned into a pEZX-MT06 dual-luciferase Target Expression Vector were provided by GeneCopoeia (*GeneCopoeia*, Rockville, MD, *USA)*. A vector without cMYC 3’UTR (GeneCopoeia) was used as experimental control. To mutate the ARE regions in the cMYC 3’UTR (**Table S7**), the sequence ARE regions was replaced as indicated (**Figure 7, Table S7**). OVCAR3 cells with stable FXR1 overexpression were transiently transfected with luciferase vectors (a luciferase vector containing the full-length (FL) cMYC 3’UTR and luciferase vectors containing the mutants (M1-M4)) with Lipofectamine 2000 Reagent (Thermo Fisher Scientific). After 48h, the luciferase activity was measured by using the Dual-Luciferase Reporter Assay System (GeneCopoeia, USA). Data were presented as the ratios between the Firefly and Renilla luminescence activities.

### Copy number variation analysis

Genome-wide copy number variation data of ovarian cancer and breast cancer were downloaded from Broad GDAC Firehose (https://gdac.broadinstitute.org/). GISTIC2 was used to identify genomic regions that are significantly gained or lost across a set of tumors. A ’+2/-2’ indicates that the sample had high-level copy number amplification or deletion. We defined the copy number amplification/deletion frequency as the number of patients with copy number amplification/deletion divided the total number of patients sequenced. The copy number of all patients across 3q26.33 was plotted as heat map using the R package heatmap (https://cran.r-project.org/package=pheatmap).

### Gene set enrichment analysis (GSEA)

Gene expression data of ovarian cancer, breast cancer, LUAD and LUSC were obtained from TCGA project. The expressions of protein coding genes were measured by fragments per kilobase of exon model per million reads mapped (FPKM). The expression of each gene was log2 transformed and then we calculated the Pearson Correlation Coefficient (PCC) between the expression of all other genes and FXR1. All genes were ranked based on PCC and then subjected to GSEA analysis [40]. The enrichment score (ES) was calculated for each functional set, which reflects the degree to which a gene set is overrepresented at the top or bottom of the ranked list of genes. Moreover, the normalized enrichment score (NES) was calculated based on 1000 permutations. Here, the cancer hallmark gene sets from MSigDB were considered for the names and the gene sets with false discovery rate <0.001 were considered as a selection criteria [41].

### Clinical data analysis using cBioPortal

TCGA data datasets were first analyzed using cBioPortal (http://www.cbioportal.org/). The segmented data for all the samples were downloaded from the TCGA Firehose (http://firebrowse.org, version: 20160128), followed by standard GISTIC2 (Genomic Identification of Significant Targets in Cancer, version 2) analysis using Firehose-suggested parameters. We evaluated the protein expression of FXR1 in multiple cancers by Clinical Proteomic Tumor Analysis Consortium (CPTAC) analysis using UALCAN data portal (http://ualcan.path.uab.edu/) [42].

### Statistical analysis

In most cases, data obtained from three or four biological replicates were analyzed, unless indicated otherwise in the Figureure legends. Statistical significance defined as a P value < 0.05 or < 0.01 was determined by unpaired Student’s t-test. Comparisons in multiple groups were analyzed with one-way or two-way analysis of variance (ANOVA), which is mentioned in the respective Figureure legends. Data are presented as the mean ± standard error (SEM) as indicated in the Figureure legends. For the analysis of correlation co-efficient, Pearson’s correlation coefficients (r) were calculated. Heatmaps were prepared with heatmapper software http://www.heatmapper.ca/. Graphpad Prism 7 (GraphPad, San Diego, CA) was used to perform statistical analysis and p-value determinations.

## Author contributions

P.C.R. conceived the study, generated hypotheses, and designed the experiments. J.G. performed most of the experiments, including cell cultures, animal experiments, qPCR, microscopy, immunoblots, statistical analyses, prepared Figures and the draft of the manuscript. Y.L. and S-W.T. performed all the bioinformatics and computational analysis for this study. D.P., A.G., P.G., and I.P.K assisted on animal experiments, animal imaging or in vitro experiments. S.P. and P.C.R. designed the animal experiments. S.P. provided scientific feedback and assisted with manuscript preparation. Y.S., and H.R., assisted on IHC scanning and pathology analysis consultations. H.R., J.S.R, R.R. and M.D. edited the manuscript and provided comments. C.G., and M.D. assisted with polysome fractionation and translational experiments. P.C.R. provided scientific direction, established collaborations, prepared the manuscript with J.G., and allocated funding for the work.

## Acknowledgements

This work was supported in part by funding from the Ovarian Cancer Research Fund Alliance (OCRFA; P.C.R., S.P.), DoD Breast Cancer Research Program (W81XWH-18-1-0024; P.C.-R), the Women’s Health Research Program (WHRP) at MCW (P.C.-R, S.P.), MCW Cancer Center Institutional Research Grants from the American Cancer Society (16-183-31 to P.C.-R., 14-247-29 to Y.S.), and research funds from MCW Cancer Center. Critical support was also received from the NCI R01CA229907 to P.C.-R., R01CA188575 to H.R., NIGMS 1R01GM124183 to M.D. and the National Natural Science Foundation of China (31970646 to Y. L.). Authors acknowledge Dr. Dinah Singer, Center for Cancer Research, National Cancer Institute for her guidance and feedback on translational assays.

