## Supplementary Text and Figures for "RNA-binding protein FXR1 drives cMYC translation by mRNA circularization through eIF4F recruitment in ovarian cancer"

### SUPPLEMENTAL DATA

### Fig S1

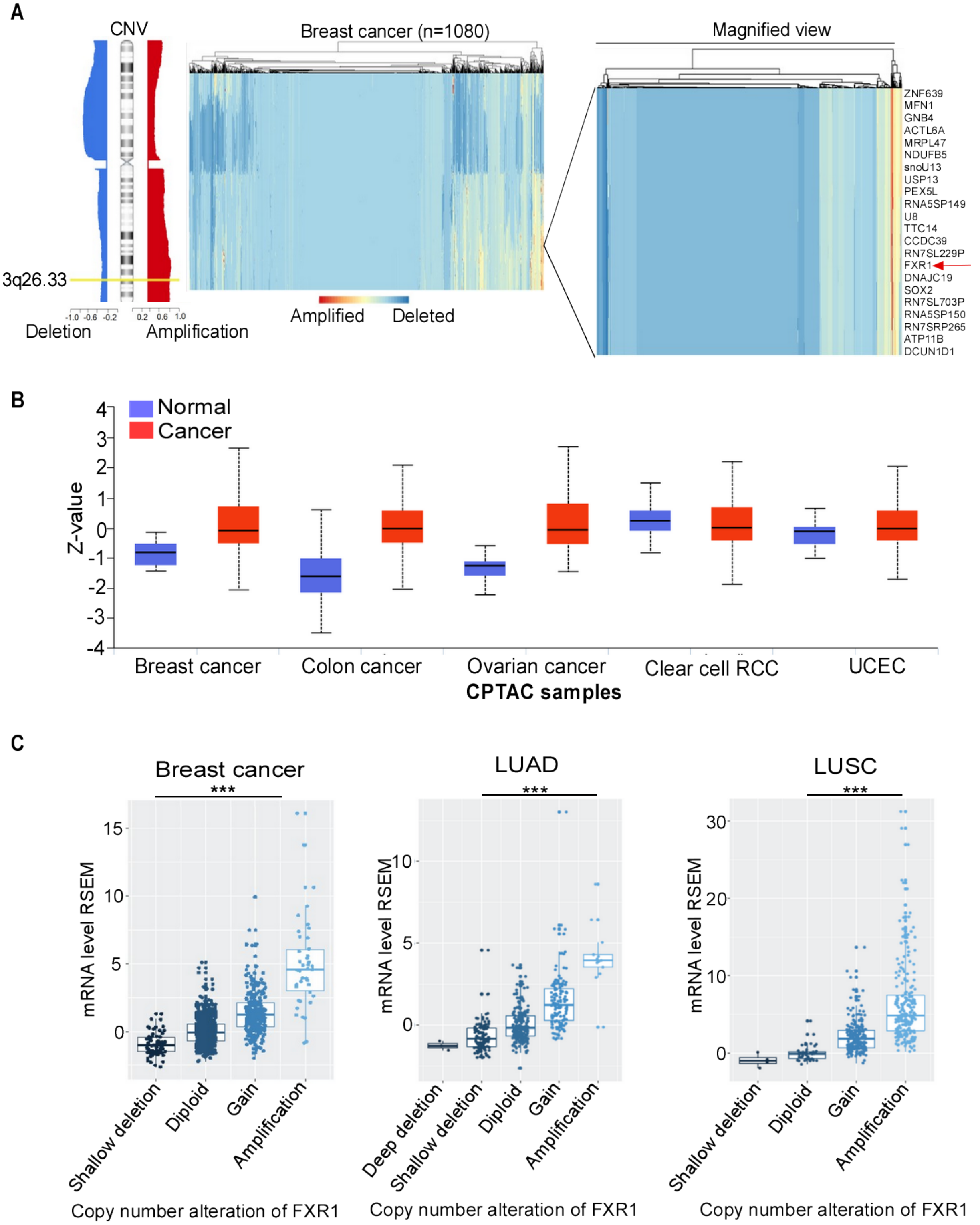

D

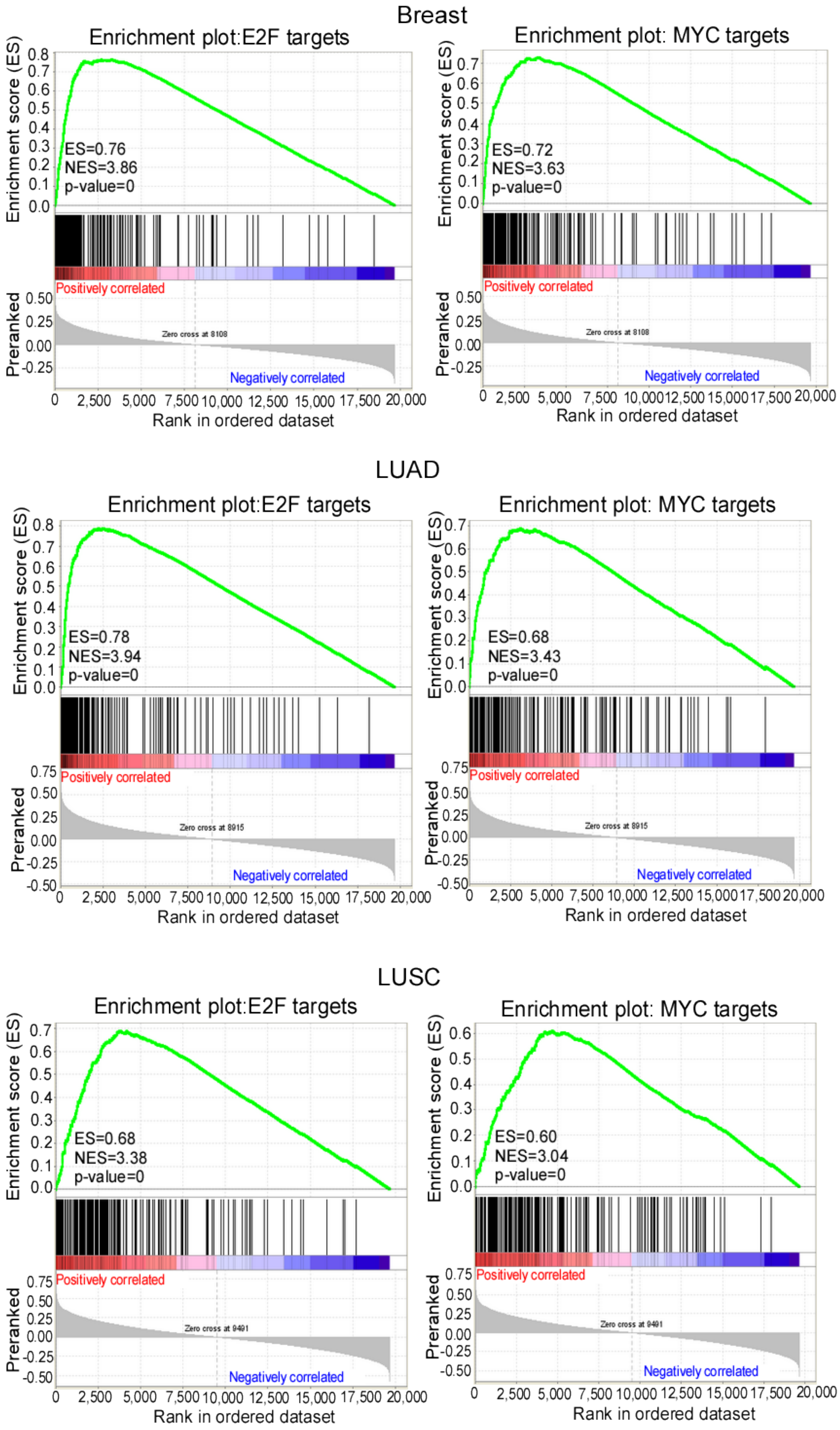
Breast
Enrichment plot:E2F targets
ES=0.76
NES=3.86
p-value=0
Enrichment plot: MYC targets
ES=0.72
NES=3.63
p-value=0
LUAD
Enrichment plot:E2F targets
ES=0.78
NES=3.94
p-value=0
Enrichment plot: MYC targets
ES=0.68
NES=3.43
p-value=0
LUSC
Enrichment plot:E2F targets
ES=0.68
NES=3.38
p-value=0
Enrichment plot: MYC targets
ES=0.60
NES=3.04
p-value=0
Enrichment score (ES)
Preranked
Positively correlated
Negatively correlated
Rank in ordered dataset

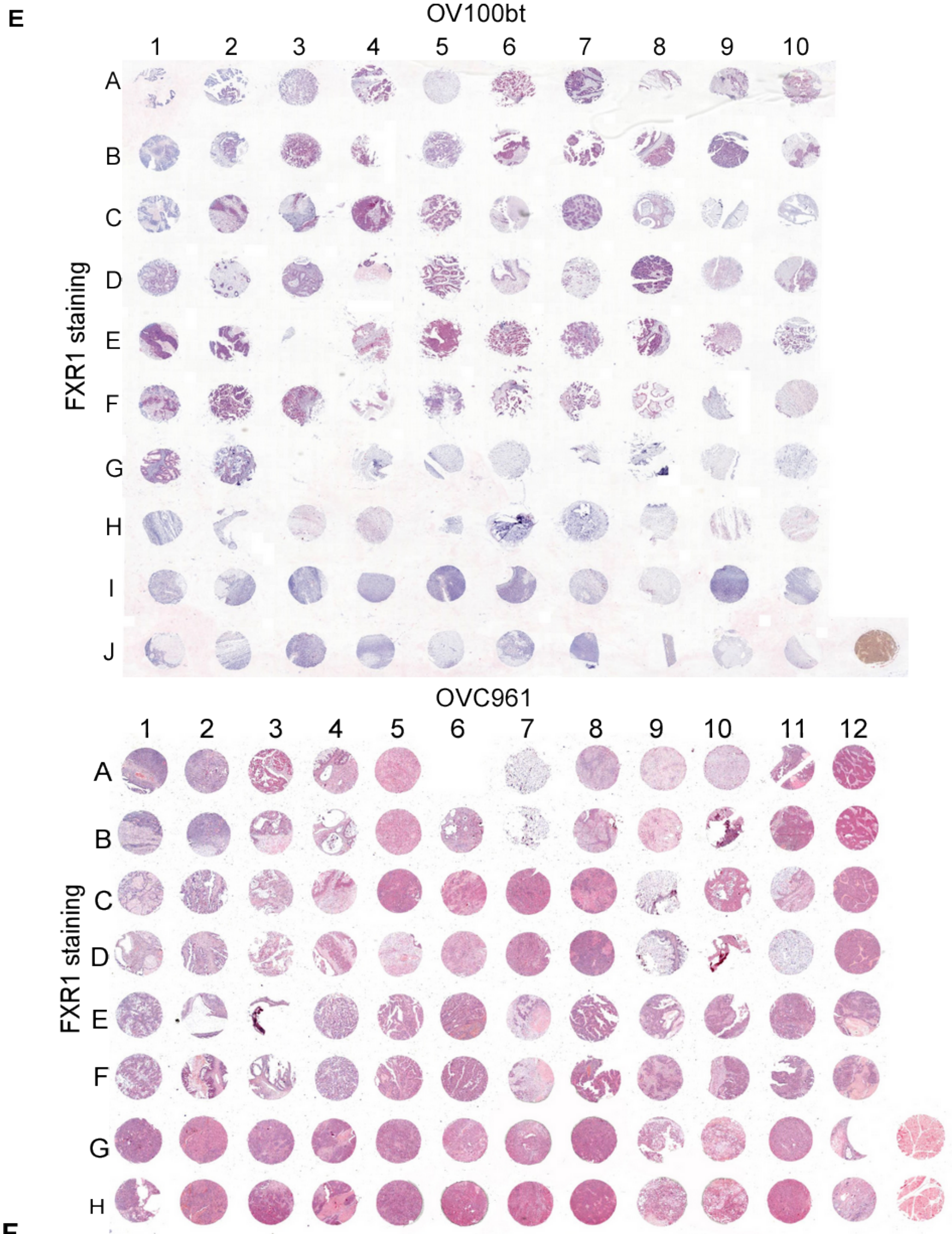
E
OV100bt
1 2 3 4 5 6 7 8 9 10
A B C D E F G H I J
FXR1 staining
OVC961
1 2 3 4 5 6 7 8 9 10 11 12
A B C D E F G H
FXR1 staining

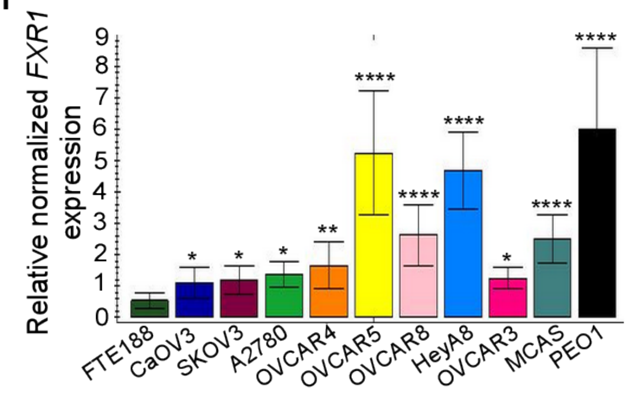
F
Relative normalized FXR1 expression
0 1 2 3 4 5 6 7 8 9
*
*
*
**
****
****
****
*
****
****
FTE188
CaOV3
SKOV3
A2780
OVCAR4
OVCAR5
OVCAR8
HeyA8
OVCAR3
MCAS
PEO1

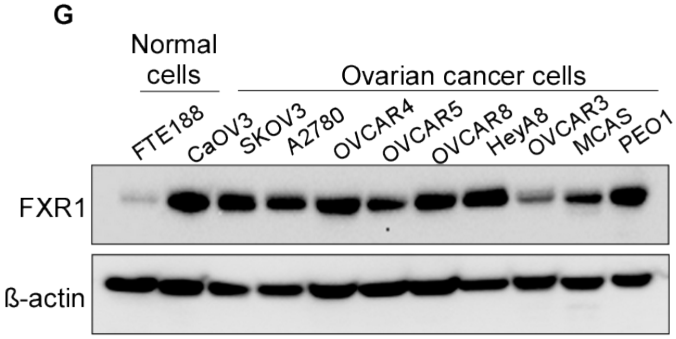
G
Normal cells
Ovarian cancer cells
FTE188
CaOV3
SKOV3
A2780
OVCAR4
OVCAR5
OVCAR8
HeyA8
OVCAR3
MCAS
PEO1
FXR1
ß-actin

**Fig S1: FXR1 is positively correlated with cancer progression. (A)** Heatmap of chromosome 3q26.3 locus amplification in TCGA breast cancer patient dataset (n=1080). Representative genes in this amplicon is magnified, where red arrow indicate the position of FXR1. **(B)** Boxplot showing relative expression of FXR1 protein across different cancers retrieved from the Clinical Proteomic Tumor Analysis Consortium (CPTAC) dataset assessed in UALCAN web portal. Clear cell renal cell carcinoma (RCC), UCEC (Uterine corpus endometrial carcinoma). **(C)** Box plots shows the association of the copy number alterations of *FXR1* mRNA. These considered with respect to the median expression of all data in the TCGA breast cancer, lung adenocarcinoma (LUAD) and lung squamous cell carcinoma (LUSC) data set. ***$p<0.001$ compared between adjacent groups. **(D)** GSEA analysis demonstrates the enrichment score of indicated functional annotation marks based on the correlation between expression of all genes and FXR1 in the TCGA breast cancer, LUAD and LUSC samples. ES: enrichment score, NES: normalized enrichment score. **(E)** Overview of immunohistochemical staining for FXR1 in the whole slide (x0.5 magnification) of human ovarian cancer tissue microarrays. Two tissue microarrays of ovarian cancer were used in this study: Cat#OV100b and Cat#OVC961. The slides were stained for FXR1 and scanned using an Aperio Scan Scope (Aperio Technologies). **(F)** Quantitative bar graph for *FXR1* mRNA expression in human normal epithelial fallopian tube cells (FTE188) and human ovarian cancer cell lines. *GAPDH* was used as a loading control. Error bars indicate SEM.* $p<0.05$, **$p<0.01$, ****$p<0.0001$ compared to FTE188 by one-way Anova test. **(G)** Western blot analysis of FXR1 levels in human normal epithelial fallopian cells and human ovarian cancer cells. β-actin was used as loading control.

### Fig S2

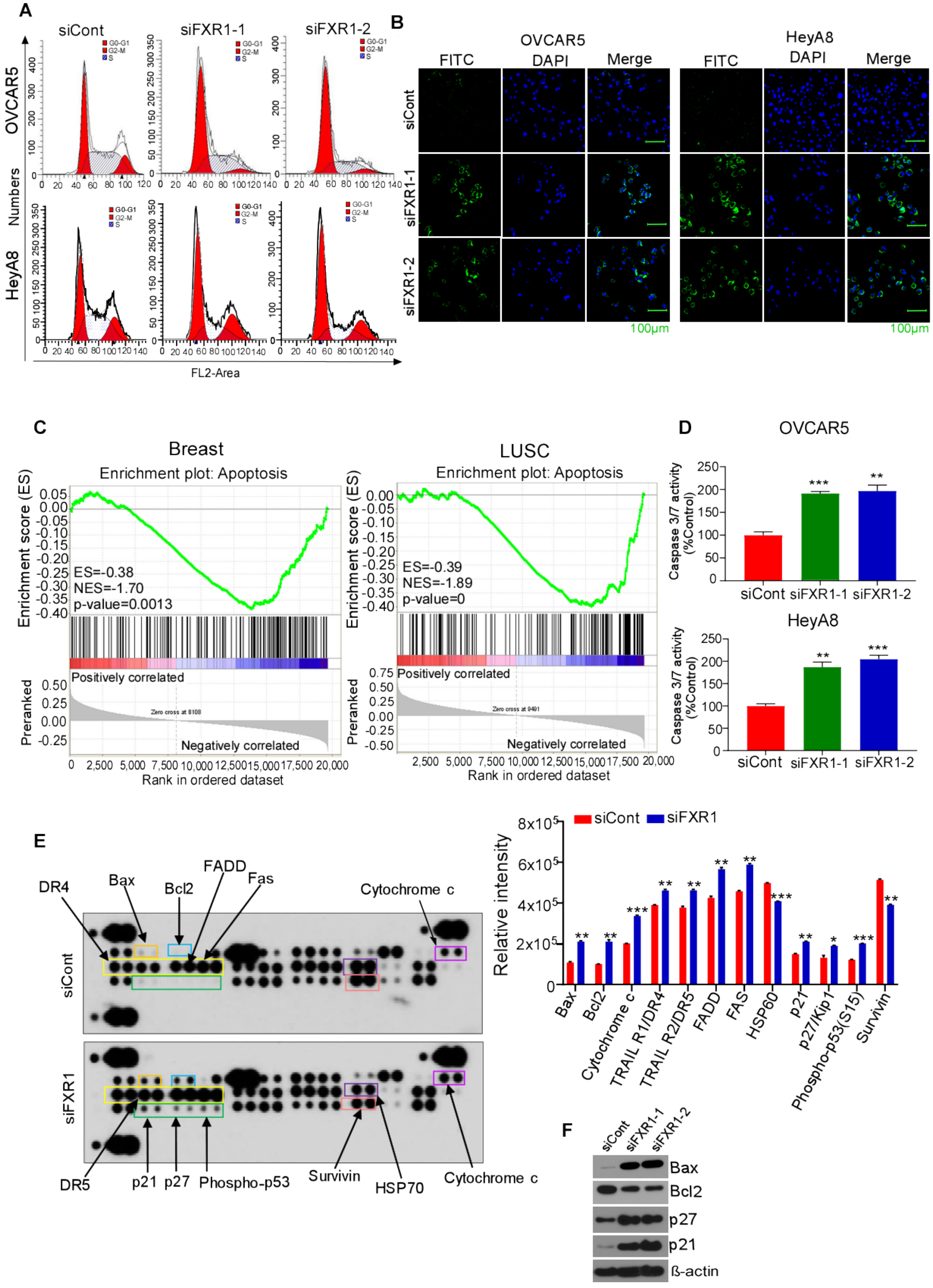

A
siCont
siFXR1-1
siFXR1-2
OVCAR5
HeyA8
Numbers
FL2-Area
G0-G1
G2-M
S
B
OVCAR5
HeyA8
FITC
DAPI
Merge
siCont
siFXR1-1
siFXR1-2
100μm
C
Breast
LUSC
Enrichment plot: Apoptosis
Enrichment score (ES)
ES=-0.38
NES=-1.70
p-value=0.0013
ES=-0.39
NES=-1.89
p-value=0
Positively correlated
Negatively correlated
Preranked
Rank in ordered dataset
D
OVCAR5
HeyA8
Caspase 3/7 activity (%Control)
siCont
siFXR1-1
siFXR1-2
E
DR4
Bax
Bcl2
FADD
Fas
Cytochrome c
siCont
siFXR1
DR5
p21
p27
Phospho-p53
Survivin
HSP70
Cytochrome c
Relative intensity
siCont
siFXR1
Bax
Bcl2
Cytochrome c
TRAIL R1/DR4
TRAIL R2/DR5
FADD
FAS
HSP60
p21
p27/Kip1
Phospho-p53(S15)
Survivin
F
siCont
siFXR1-1
siFXR1-2
Bax
Bcl2
p27
p21
ß-actin

**Fig S2: Knockdown of FXR1 induced G1 phase arrest and apoptosis in ovarian cancer cells. (A)** Representative histograms depicting cell cycle distribution in OVCAR5 and HeyA8 cells were transfected with control siRNA (siCont) or FXR1 siRNA (siFXR1-1 and siFXR1-2) for 48h. **(B)** Representative confocal microscopy images (x20 magnification) of OVCAR5 and HeyA8 cells were stained with Live-or-Dye™ 488/515 after treatment with siCont and FXR1 siRNAs for 48h. Nuclei were counterstained with DAPI. **(C)** GSEA analysis demonstrates the enrichment score of indicated functional annotation marks based on FXR1 expression in the TCGA breast and LUSC cancer samples. ES: enrichment score, NES: normalized enrichment score. **(D)** Quantitative bar graph showing increase in caspase 3/7 activity after siRNAs knockdown of FXR1 in ovarian cancer cells. Luminescence intensity was measured using plate reader. Error bars indicate SEM. **$p<0.01$, ***$p<0.001$ compared to siCont by Student's t-test. **(E)** HeyA8 cells were transfected with siCont or siFXR1 siRNAs for 48h and then lysates were prepared and apoptosis protein array (R and D Systems) was performed. Significantly altered proteins were marked in boxes (left). Densitometry quantification of proteins (right) levels marked in boxes from protein array. Error bars indicate SEM.* $p<0.05$, **$p<0.01$, ***$p<0.001$ compared to siCont by Student's t-test. **(F)** Cell lysates from E were blotted for the indicated proteins. β-actin was used as loading controls.

Fig S3

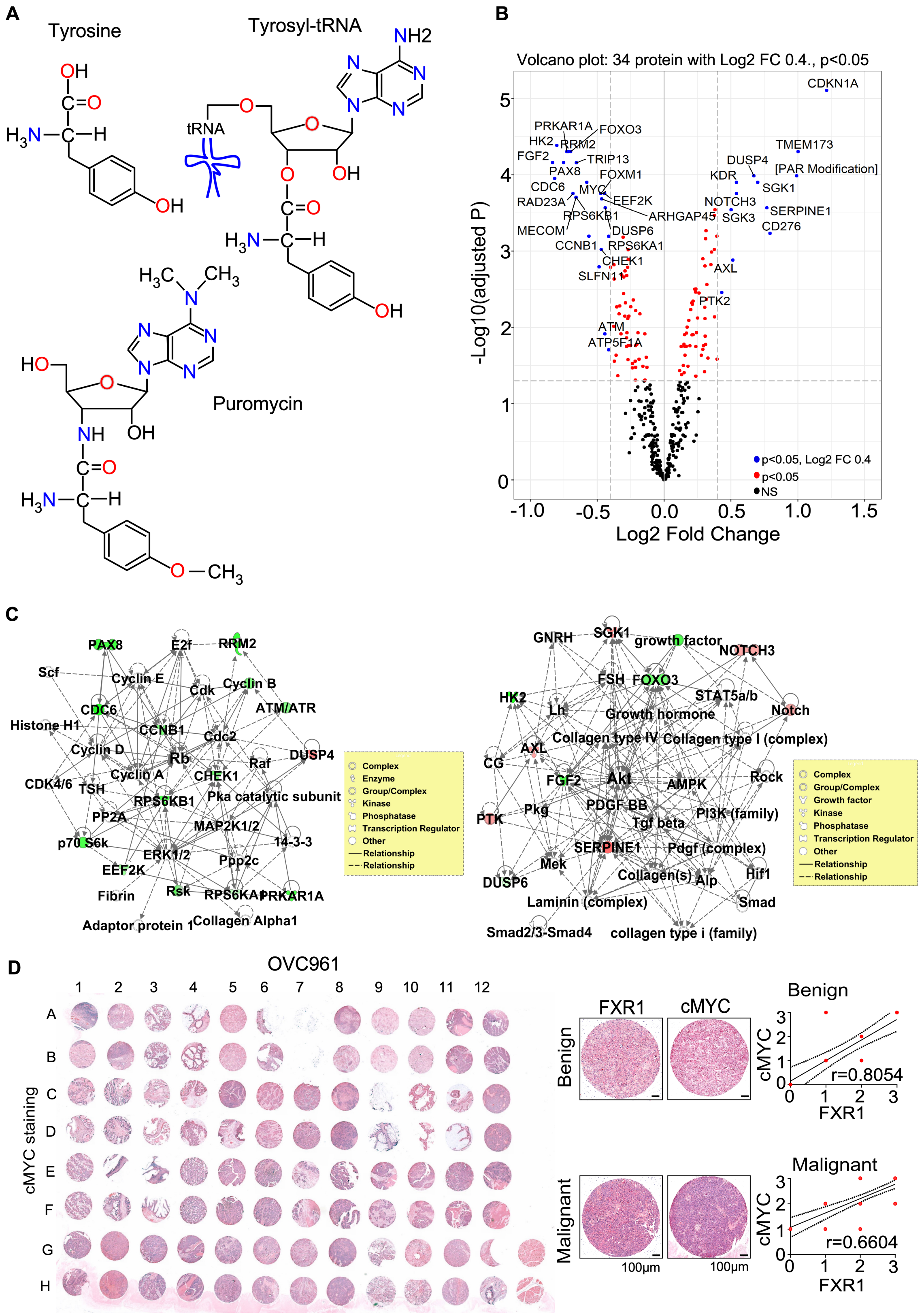
A
Tyrosine
Tyrosyl-tRNA
tRNA
Puromycin
B
Volcano plot: 34 protein with Log2 FC 0.4., p<0.05
-Log10(adjusted P)
Log2 Fold Change
p<0.05, Log2 FC 0.4
p<0.05
NS
CDKN1A
TMEM173
DUSP4
[PAR Modification]
KDR
SGK1
NOTCH3
SERPINE1
SGK3
CD276
AXL
PTK2
PRKAR1A
FOXO3
HK2
RRM2
FGF2
TRIP13
PAX8
FOXM1
CDC6
MYC
RAD23A
EEF2K
RPS6KB1
ARHGAP45
MECOM
DUSP6
CCNB1
RPS6KA1
CHEK1
SLFN11
ATM
ATP5F1A
C
Complex
Enzyme
Group/Complex
Kinase
Phosphatase
Transcription Regulator
Other
Relationship
Relationship
Growth factor
D
OVC961
cMYC staining
FXR1
cMYC
Benign
Malignant
r=0.8054
r=0.6604
100μm
100μm

**Fig S3: FXR1 effects translation of downstream target mRNAs including cMYC. (A)** Molecular structures of tyrosine, tyrosyl-tRNA, and puromycin. Aminoacyl-tRNA contain a hydrolyzable ester bond between their tRNA ribose moiety and the attached amino acid, whereas puromycin has a nonhydrolyzable amide bond in the equivalent position. **(B)** Volcano plot of the significantly differentially expressed proteins (n=34) from Reverse phase protein array (RPPA) data. Volcano plots showing Log2 fold changes (FC) against Log10 p-values of differentially expressed proteins in HeyA8 cells treated with siFXR1 for 48h compared to siCont. Blue dots represent proteins, which are significantly down-regulated or up-regulated expression with $p<0.05$ with Log2 FC 0.4, while red dots represent proteins with significant expression $p<0.05$ and black spots represents non-significant (NS) changes. **(C)** Top two and top three gene networks plotted by Ingenuity Pathway Analysis (IPA) for differentially expressed proteins (n= 34, Log2 fold change) from RPPA as described in **Fig. 3F**. Genes are represented by nodes with their shape representing the type of molecule/functional class, and the relationship between the nodes are indicated by edges. Green highlighted proteins indicate downregulated and red indicate upregulated. Solid lines indicate direct protein–protein interactions, dashed lines indicate potential protein–protein interaction, and arrows imply directionality of protein–protein regulation. A p-value < 0.05 by Fisher's exact t test performed to determine significance. **(D)** Immunohistochemical staining for cMYC in the whole slide of human ovarian cancer tissue microarray slide (left); Magnification: 0.5x. Representative images of benign and malignant ovarian tumor tissue of cMYC from **D** and representative images of cMYC from S1E presented for correlation coefficient analysis Magnification: 10x (middle). Dot graph with regression line plotted based on IHC score in benign and malignant samples; r = Pearson's coefficient value (right).

**Fig S4**

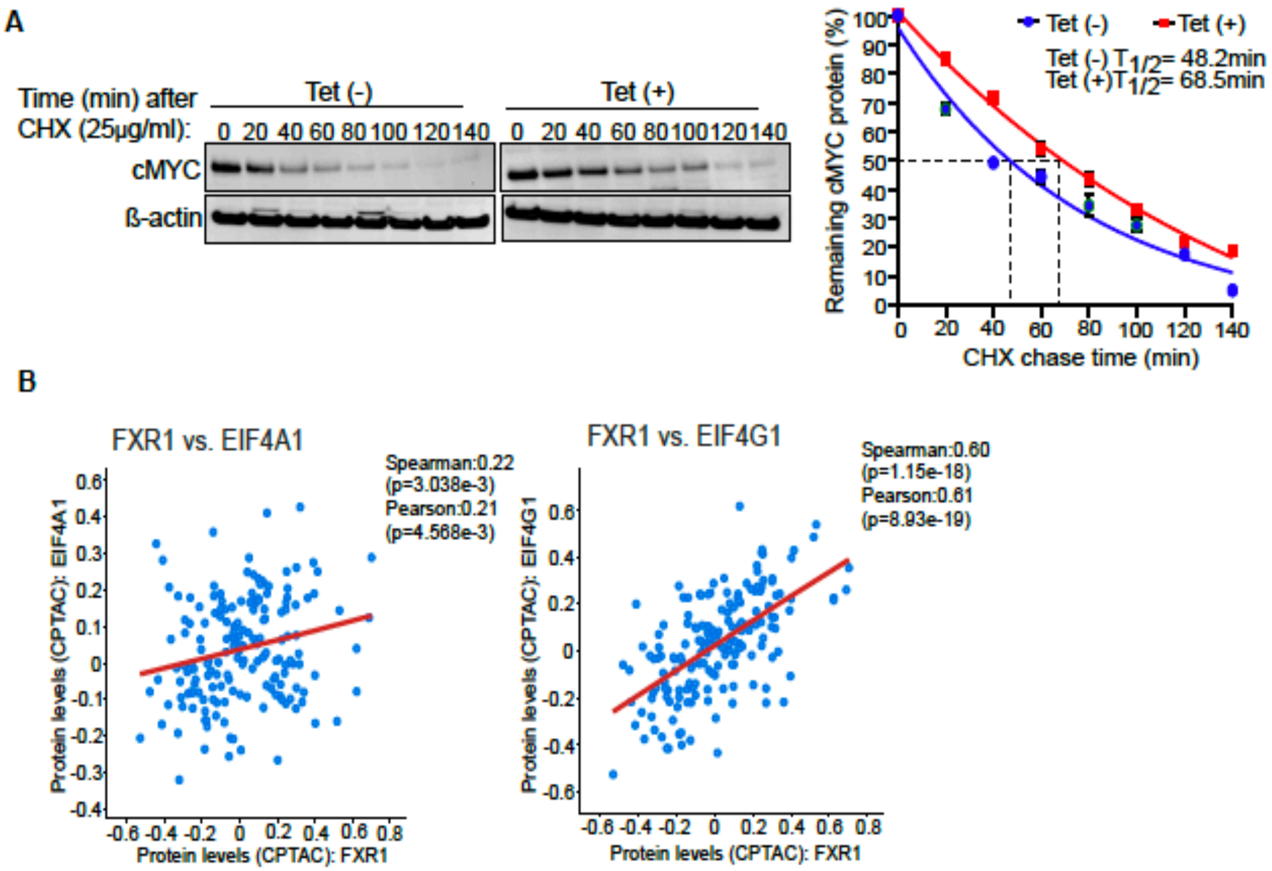


**Fig S4: FXR1 maintains cMYC stability and interacts with eukaryotic translation initiation complex. (A)** Immunoblot was performed using the lysates prepared from tetracycline inducible FXR1 overexpressing HeyA8 cells (Tet (+)), followed by the treatment with cycloheximide (25μg/ml) for indicated time points (left). Densitometric quantification for phase decay of cMYC protein bands from the blots performed using image-J software (right). **(B)** Scatter plot with regression line present correlation between FXR1 with eIF4G1, and eIF4A1, proteins in TCGA ovarian cancer dataset. All correlation values were calculated by Spearman and Pearson correlation analysis.

### Fig S5

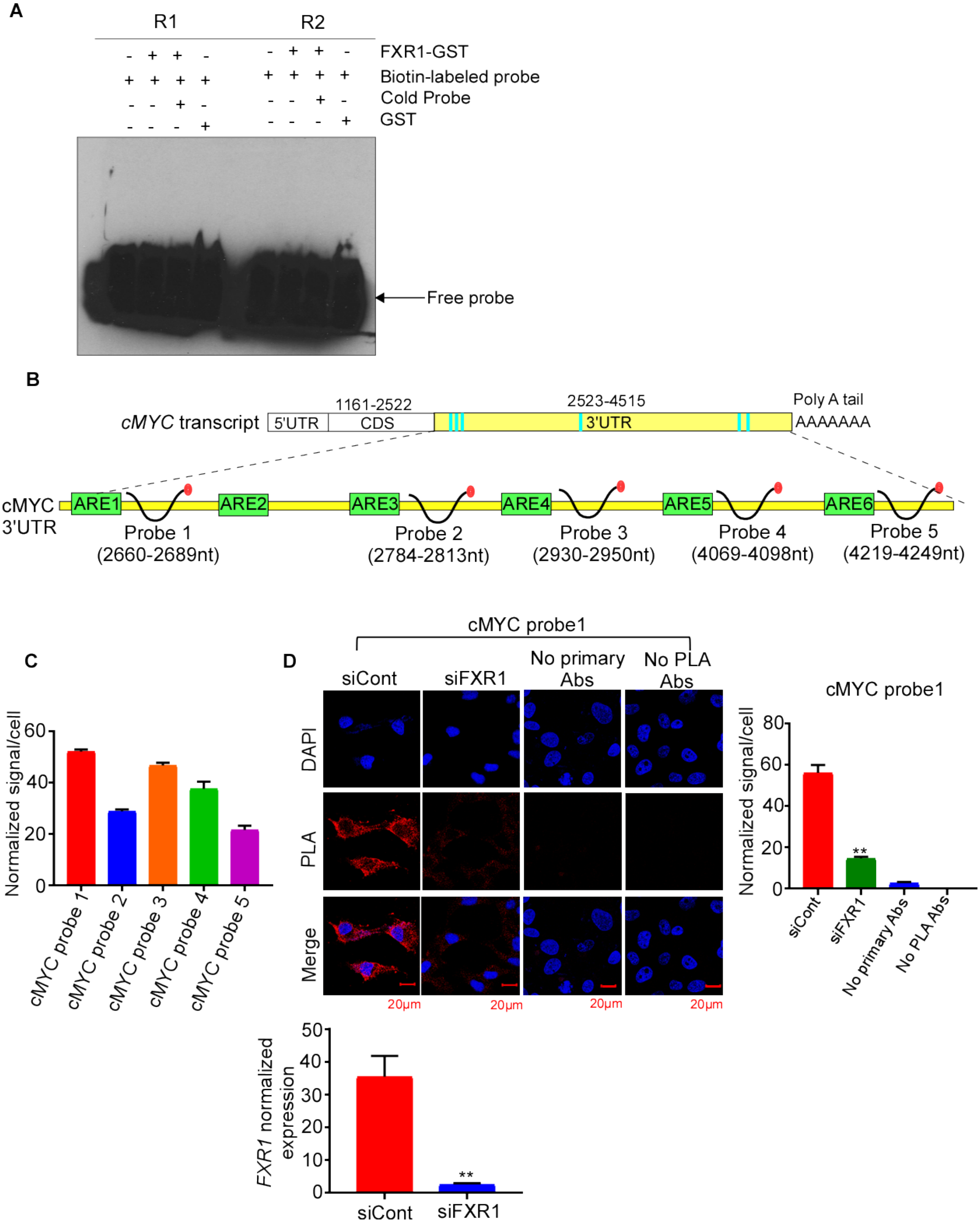
A
R1
R2
FXR1-GST
Biotin-labeled probe
Cold Probe
GST
Free probe
B
cMYC transcript
5'UTR
1161-2522
CDS
2523-4515
3'UTR
Poly A tail
AAAAAAA
cMYC 3'UTR
ARE1
ARE2
ARE3
ARE4
ARE5
ARE6
Probe 1 (2660-2689nt)
Probe 2 (2784-2813nt)
Probe 3 (2930-2950nt)
Probe 4 (4069-4098nt)
Probe 5 (4219-4249nt)
C
Normalized signal/cell
cMYC probe 1
cMYC probe 2
cMYC probe 3
cMYC probe 4
cMYC probe 5
D
cMYC probe1
siCont
siFXR1
No primary Abs
No PLA Abs
DAPI
PLA
Merge
20μm
Normalized signal/cell
cMYC probe1
**
FXR1 normalized expression
siCont
siFXR1
**

**Fig S5: FXR1 binds to the 3′UTRs within *cMYC* transcript. (A)** Biotinylated random RNA probes (R1(3108-3122nt) and R2(3809-3823nt)) containing 15-mer of human cMYC ARE were incubated with human FXR1-GST recombinant protein or with GST alone overnight. For competition assay, biotinylated RNA probes were mixed with or without 200-fold of unlabeled cMYC probes and incubated overnight. Reaction mixtures were then resolved on 6% gel and blotted to nylon membrane and then developed by chemiluminescence assay. Arrow marks indicate the position of free probe and shift in the mobility due to FXR1 binding. **(B)** Schematic representation of *cMYC* transcript and the location of probes in the 3'UTR used for PLA. Green color indicates the position AREs and black curve with red-tail indicates biotinylated RNA probes and its proximal position to ARE. **(C)** Quantitative graph from Fig. 7E of PLA signal for FXR1 protein with *cMYC* mRNA per cell. Signals per cell were counted and normalized to the total area of the cell using LSM510 software (Carl Zeiss, Inc). Error bars represent SEM from quantifications of three cells from three different experiments are shown. **(D)** Representative confocal microscopy images (upper panel) of HeyA8 cells were transfected with siCont and siFXR1 for 48h, no primary antibody (Abs) control and, no PLA Abs control were shown Quantitative graph of *FXR1* mRNA expression (lower panel) in HeyA8 cells were treated with siCont and siFXR1 treatment for 48h. **$p<0.01$ compared to siCont by Student's t-test. Signal per cell were counted and normalized to the total area of the cell using software LSM510 (Carl Zeiss, Inc) and represented as histograms. Error bars represent SEM from quantifications of at least three cells from three different experiments are shown. **$p<0.01$ compared to siCont by Student's t-test; magnification=63x.

**Videos related to Fig 5.**

**Video S1-S2:** Positron emission tomography (PET) recording showing tumor growth and metastasis in a representative control mouse (Dox (-)) and Dox inducible FXR1 mouse (Dox (+)) at day 28, respectively.

**Table S1, related to method section of immunohistochemistry**

**Human ovarian cancer TMA (OV100b) immnuostained for FXR1 with pathological case summary for patients illustrated in Fig 1.**

| Position | No. | Age | Organ | Pathology | TNM | Grade | Stage | Type | Score |
|---|---|---|---|---|---|---|---|---|---|
| A1 | 1 | 65 | Ovary | Serous papillary cystadenocarcinoma | T1N0M0 | 1 | I | Malignant | 1+ |
| A2 | 2 | 38 | Ovary | Serous papillary cystadenocarcinoma | T3cN1M1 | 1 | IIIc | Malignant | 1+ |
| A3 | 3 | 51 | Ovary | Serous papillary cystadenocarcinoma | T3cN1M0 | 1 | IIIc | Malignant | 1+ |
| A4 | 4 | 22 | Ovary | Serous papillary cystadenocarcinoma | T2bN0M0 | 1 | IIb | Malignant | 2+ |
| A5 | 5 | 48 | Ovary | Serous papillary cystadenocarcinoma | T1N0M0 | 1 | I | Malignant | Trace |
| A6 | 6 | 26 | Ovary | Serous papillary cystadenocarcinoma | T3cN1M0 | 1 | IIIc | Malignant | 2+ |
| A7 | 7 | 25 | Ovary | Serous papillary cystadenocarcinoma | T1N0M0 | 1 | I | Malignant | 3+ |
| A8 | 8 | 50 | Ovary | Serous papillary cystadenocarcinoma | T2N0M0 | 1 | II | Malignant | 2+ |
| A9 | 9 | 26 | Ovary | Serous papillary cystadenocarcinoma | T1cN0M0 | 1 | Ic | Malignant | 2+ |
| A10 | 10 | 47 | Ovary | Serous papillary cystadenocarcinoma | T1N0M0 | 1 | I | Malignant | 2+ |
| B1 | 11 | 58 | Ovary | Serous papillary adenocarcinoma with necrosis | T1N0M0 | 2 | I | Malignant | Negative |
| B2 | 12 | 57 | Ovary | Serous papillary cystadenocarcinoma | T1cN0M0 | 2 | Ic | Malignant | 1+ |
| B3 | 13 | 51 | Ovary | Serous papillary adenocarcinoma | T1aN0M0 | 2 | Ia | Malignant | 3+ |
| B4 | 14 | 52 | Ovary | Serous papillary cystadenocarcinoma | T2N0M0 | 2 | II | Malignant | 2+ |
| B5 | 15 | 54 | Ovary | Serous papillary adenocarcinoma | T3cN1M0 | 2 | IIIc | Malignant | 1+ |
| B6 | 16 | 33 | Ovary | Serous papillary adenocarcinoma | T1N0M0 | 2 | I | Malignant | 2+ |
| B7 | 17 | 56 | Ovary | Serous papillary adenocarcinoma | T2N0M0 | 3 | II | Malignant | 3+ |
| B8 | 18 | 41 | Ovary | Serous papillary adenocarcinoma | T1N0M0 | 2 | I | Malignant | 3+ |
| B9 | 19 | 46 | Ovary | Serous papillary adenocarcinoma | T3aN0M0 | 2 | IIIc | Malignant | 3+ |
| B10 | 20 | 46 | Ovary | Serous papillary adenocarcinoma | T2cN1M0 | 2 | IIIc | Malignant | 2+ |

| | | | | | | | | | |
|---|---|---|---|---|---|---|---|---|---|
| C1 | 21 | 57 | Ovary | Serous adenocarcinoma | T3cN1M0 | 2 | IIIc | Malignant | Negative |
| C2 | 22 | 75 | Ovary | Serous adenocarcinoma | T2N0M0 | 2--3 | II | Malignant | 3+ |
| C3 | 23 | 54 | Ovary | Serous adenocarcinoma | T3cN1M0 | 3 | IIIc | Malignant | 2+ |
| C4 | 24 | 49 | Ovary | Serous adenocarcinoma | T2N0M0 | 3 | II | Malignant | 3+ |
| C5 | 25 | 50 | Ovary | Serous papillary adenocarcinoma | T1N0M0 | 2 | I | Malignant | 3+ |
| C6 | 26 | 52 | Ovary | Serous adenocarcinoma | T2N0M0 | 3 | II | Malignant | Trace |
| C7 | 27 | 47 | Ovary | Serous adenocarcinoma | T3cN1M0 | 3 | IIIc | Malignant | 3+ |
| C8 | 28 | 34 | Ovary | Mucinous adenocarcinoma | T1bN0M0 | 2 | Ib | Malignant | 1+ |
| C9 | 29 | 63 | Ovary | Mucinous adenocarcinoma | T1aN0M0 | 1 | Ia | Malignant | Negative |
| C10 | 30 | 69 | Ovary | Mucinous adenocarcinoma | T1bN0M0 | 1 | Ib | Malignant | Negative |
| D1 | 31 | 46 | Ovary | Endometrioid adenocarcinoma | T2N0M0 | 1--2 | II | Malignant | 1+ |
| D2 | 32 | 47 | Ovary | Endometrioid adenocarcinoma | T2aN0M0 | 1--2 | IIa | Malignant | 1+ |
| D3 | 33 | 54 | Ovary | Endometrioid adenocarcinoma | T1bN0M0 | 1--2 | Ib | Malignant | 2+ |
| D4 | 34 | 65 | Ovary | Adenocarcinoma (sparse) | T1cN0M0 | - | Ic | Malignant | Trace |
| D5 | 35 | 55 | Ovary | Endometrioid adenocarcinoma | T1N0M0 | 2 | I | Malignant | 3+ |
| D6 | 36 | 54 | Ovary | Endometrioid adenocarcinoma | T1bN0M0 | 1 | Ib | Malignant | 2+ |
| D7 | 37 | 43 | Ovary | Endometrioid adenocarcinoma | T1cN0M0 | 1 | Ic | Malignant | 1+ |
| D8 | 38 | 55 | Ovary | Endometrioid adenocarcinoma | T1N0M0 | 1 | I | Malignant | 3+ |
| D9 | 39 | 53 | Ovary | Endometrioid adenocarcinoma | T2aN0M0 | 3 | IIa | Malignant | 1+ |
| D10 | 40 | 50 | Ovary | Endometrioid adenocarcinoma | T3bN1M0 | 2 | IIIc | Malignant | 1+ |
| E1 | 41 | 51 | Ovary | Transitional cell carcinoma | T1bN0M0 | 2 | Ib | Malignant | 3+ |
| E2 | 42 | 39 | Ovary | Transitional cell carcinoma | T1aN0M0 | 2 | Ia | Malignant | 3+ |
| E3 | 43 | 38 | Ovary | Transitional cell carcinoma | T1N0M0 | 2--3 | I | Malignant | Negative |
| E4 | 44 | 66 | Ovary | Transitional cell carcinoma | T1aN0M0 | 3 | Ia | Malignant | 2+ |
| E5 | 45 | 53 | Ovary | Transitional cell carcinoma | T1N0M0 | 2--3 | I | Malignant | 3+ |
| E6 | 46 | 47 | Mesentery | Metastatic serous papillary cystadenocarcinoma from ovary | - | 1 | - | Metastasis | 3+ |
| E7 | 47 | 57 | Epiploon | Metastatic serous papillary cystadenocarcinoma from ovary | - | 1 | - | Metastasis | 3+ |

| | | | | | | | | | |
|---|---|---|---|---|---|---|---|---|---|
| E8 | 48 | 65 | Epiploon | Metastatic serous papillary cystadenocarcinoma with calcification from ovary | - | 1 | - | Metastasis | 3+ |
| E9 | 49 | 59 | Mesentery | Metastatic serous papillary cystadenocarcinoma from ovary | - | 2 | - | Metastasis | 2+ |
| E10 | 50 | 28 | Epiploon | Metastatic serous adenocarcinoma with calcification from ovary | - | 2 | - | Metastasis | 2+ |
| F1 | 51 | 64 | Epiploon | Metastatic serous papillary cystadenocarcinoma from ovary | - | 1 | - | Metastasis | 3+ |
| F2 | 52 | 50 | Epiploon | Metastatic serous papillary cystadenocarcinoma from ovary | - | 1 | - | Metastasis | 3+ |
| F3 | 53 | 58 | Epiploon | Metastatic adenocarcinoma from ovary | - | 2 | - | Metastasis | 3+ |
| F4 | 54 | 47 | Abdominal cavity | Metastatic adenocarcinoma from ovary | - | 2 | - | Metastasis | Trace |
| F5 | 55 | 49 | Abdominal cavity | Metastatic adenocarcinoma from ovary | - | 3 | - | Metastasis | 2+ |
| F6 | 56 | 34 | Ovary | Borderline serous papillary cystadenoma | - | - | - | Borderline | 2+ |
| F7 | 57 | 34 | Ovary | Borderline serous papillary cystadenoma | - | - | - | Borderline | 2+ |
| F8 | 58 | 28 | Ovary | Borderline serous papillary cystadenoma | - | - | - | Borderline | 2+ |
| F9 | 59 | 22 | Ovary | Borderline serous papillary cystadenoma | - | - | - | Borderline | Negative |
| F10 | 60 | 60 | Ovary | Borderline serous papillary cystadenoma | - | - | - | Borderline | Trace |
| G1 | 61 | 50 | Ovary | Borderline serous papillary cystadenoma | - | - | - | Borderline | 2+ |
| G2 | 62 | 37 | Ovary | Borderline mucinous papillary cystadenoma | - | - | - | Borderline | 2+ |

| | | | | | | | | | |
|---|---|---|---|---|---|---|---|---|---|
| G3 | 63 | 62 | Ovary | Serous cystadenoma | - | - | - | Benign | Negative |
| G4 | 64 | 70 | Ovary | Serous cystadenoma | - | - | - | Benign | Negative |
| G5 | 65 | 49 | Ovary | Serous cystadenoma | - | - | - | Benign | Negative |
| G6 | 66 | 16 | Ovary | Serous cystadenoma | - | - | - | Benign | Negative |
| G7 | 67 | 34 | Ovary | Serous cystadenoma | - | - | - | Benign | Negative |
| G8 | 68 | 22 | Ovary | Serous cystadenoma | - | - | - | Benign | Negative |
| G9 | 69 | 19 | Ovary | Mucinous cystadenoma | - | - | - | Benign | Negative |
| G10 | 70 | 17 | Ovary | Mucinous cystadenoma | - | - | - | Benign | Negative |
| H1 | 71 | 41 | Uterus | Mucinous cystadenoma | - | - | - | Benign | Negative |
| H2 | 72 | 26 | Ovary | Mucinous cystadenoma | - | - | - | Benign | Negative |
| H3 | 73 | 22 | Ovary | Mucinous cystadenoma | - | - | - | Benign | Trace |
| H4 | 74 | 38 | Ovary | Mucinous cystadenoma | - | - | - | Benign | Trace |
| H5 | 75 | 47 | Ovary | Mucinous cystadenoma | - | - | - | Benign | Negative |
| H6 | 76 | 70 | Ovary | Mucinous cystadenoma | - | - | - | Benign | Negative |
| H7 | 77 | 51 | Ovary | Mucinous cystadenoma | - | - | - | Benign | Negative |
| H8 | 78 | 29 | Ovary | Mucinous cystadenoma | - | - | - | Benign | Negative |
| H9 | 79 | 35 | Ovary | Mucinous cystadenoma | - | - | - | Benign | Trace |
| H10 | 80 | 18 | Ovary | Mucinous cystadenoma | - | - | - | Benign | Trace |
| I1 | 81 | 30 | Ovary | Cancer adjacent normal ovary tissue | - | - | - | NAT | Negative |
| I2 | 82 | 39 | Ovary | Cancer adjacent normal ovary tissue | - | - | - | NAT | Negative |
| I3 | 83 | 29 | Ovary | Cancer adjacent normal ovary tissue | - | - | - | NAT | Negative |
| I4 | 84 | 41 | Ovary | Cancer adjacent normal ovary tissue | - | - | - | NAT | Negative |
| I5 | 85 | 62 | Ovary | Cancer adjacent normal ovary tissue | - | - | - | NAT | Negative |
| I6 | 86 | 63 | Ovary | Cancer adjacent normal ovary tissue | - | - | - | NAT | Negative |
| I7 | 87 | 45 | Ovary | Cancer adjacent normal ovary tissue | - | - | - | NAT | Negative |
| I8 | 88 | 48 | Ovary | Cancer adjacent normal ovary tissue | - | - | - | NAT | Negative |
| I9 | 89 | 53 | Ovary | Cancer adjacent normal ovary tissue | - | - | - | NAT | Negative |
| I10 | 90 | 53 | Ovary | Cancer adjacent normal ovary tissue | - | - | - | NAT | Negative |
| J1 | 91 | 57 | Ovary | Cancer adjacent normal ovary tissue | - | - | - | NAT | Negative |
| J2 | 92 | 38 | Ovary | Cancer adjacent normal ovary tissue | - | - | - | NAT | Negative |
| J3 | 93 | 53 | Ovary | Cancer adjacent normal ovary tissue | - | - | - | NAT | Negative |

| | | | | | | | | | |
|---|---|---|---|---|---|---|---|---|---|
| J4 | 94 | 59 | Ovary | Cancer adjacent normal ovary tissue | - | - | - | NAT | Negative |
| J5 | 95 | 48 | Ovary | Cancer adjacent normal ovary tissue | - | - | - | NAT | Negative |
| J6 | 96 | 50 | Ovary | Cancer adjacent normal ovary tissue | - | - | - | NAT | Negative |
| J7 | 97 | 52 | Ovary | Cancer adjacent normal ovary tissue | - | - | - | NAT | Negative |
| J8 | 98 | 27 | Ovary | Normal ovary tissue | - | - | - | Normal | Negative |
| J9 | 99 | 34 | Ovary | Normal ovary tissue | - | - | - | Normal | Negative |
| J10 | 100 | 19 | Ovary | Normal ovary tissue | - | - | - | Normal | Negative |

Remark: Detailed pathology report can be obtained online: https://www.biomax.us/tissue
arrays/Ovary/OV1005b
NAT: Adjacent normal ovary tissue

**Table S2, related to method section of immunohistochemistry**

**Human ovarian cancer TMA (OVC961) immunostained for FXR1 and cMYC with pathological case summary for patients illustrated in Fig 1 and Fig S3D.**

| Position | No. | Age | Organ | Pathology | TNM | Grade | Type | Score | |
|---|---|---|---|---|---|---|---|---|---|
| | | | | | | | | FXR1 | cMYC |
| A1 | 1 | 29 | Ovary | Normal | | | normal | Trace | 1+ |
| A2 | 2 | 50 | Ovary | Normal | | | normal | Trace | 1+ |
| A3 | 3 | 51 | Ovary | Serous cystadenocarcinoma | T2N0M0 | II~III | malignant | 3+ | 2+ |
| A4 | 4 | 47 | Ovary | Mucinous cystadenoma | | | benign | 2+ | 1+ |
| A5 | 5 | 35 | Ovary | Thecoma | | | benign | 2+ | 2+ |
| A6 | 6 | 68 | Ovary | Mucinous cystadenoma | | | benign | Negative | Trace |
| A7 | 7 | 57 | Ovary | Mucinous cystadenoma | | | benign | Negative | Negative |
| A8 | 8 | 67 | Ovary | Thecoma | | | benign | 1+ | 3+ |
| A9 | 9 | 47 | Ovary | Thecoma | | | benign | 1+ | 1+ |
| A10 | 10 | 55 | Ovary | Thecoma | | | benign | 1+ | 1+ |
| A11 | 11 | 43 | Ovary | Granulosa cell tumor | | | benign | 3+ | 3+ |
| A12 | 12 | 52 | Ovary | Granulosa cell tumor | | | benign | 3+ | 3+ |
| B1 | 13 | 29 | Ovary | Normal | | | normal | Trace | 1+ |
| B2 | 14 | 50 | Ovary | Normal | | | normal | Trace | 1+ |
| B3 | 15 | 51 | Ovary | Serous cystadenocarcinoma | T2N0M0 | II~III | benign | 2+ | 2+ |
| B4 | 16 | 47 | Ovary | Mucinous cystadenoma | | | benign | 2+ | 2+ |
| B5 | 17 | 35 | Ovary | Thecoma | | | benign | 2+ | 2+ |
| B6 | 18 | 68 | Ovary | Mucinous cystadenoma | | | benign | 2+ | 2+ |
| B7 | 19 | 57 | Ovary | Mucinous cystadenoma | | | benign | Negative | Negative |

| | | | | | | | | | |
|---|---|---|---|---|---|---|---|---|---|
| B8 | 20 | 67 | Ovary | Thecoma | | | benign | 2+ | 1+ |
| B9 | 21 | 47 | Ovary | Thecoma | | | benign | 1+ | 1+ |
| B10 | 22 | 55 | Ovary | Thecoma | | | benign | 1+ | 1+ |
| B11 | 23 | 43 | Ovary | Granulosa cell tumor | | | benign | 3+ | 3+ |
| B12 | 24 | 52 | Ovary | Granulosa cell tumor | | | benign | 3+ | 3+ |
| C1 | 25 | 33 | Ovary | Serous cystadenocarcinoma | T1N0M0 | I~II | malignant | 1+ | 2+ |
| C2 | 26 | 34 | Ovary | Serous cystadenocarcinoma | T2N0M0 | II | malignant | 1+ | 2+ |
| C3 | 27 | 56 | Ovary | Serous cystadenocarcinoma | T1N0M0 | II | malignant | 2+ | 2+ |
| C4 | 28 | 45 | Ovary | Serous cystadenocarcinoma | T1N0M0 | II | malignant | 2+ | 2+ |
| C5 | 29 | 43 | Ovary | Serous cystadenocarcinoma | T1N0M0 | III | malignant | 3+ | 3+ |
| C6 | 30 | 51 | Ovary | Serous cystadenocarcinoma | T3N1M0 | III | malignant | 3+ | 3+ |
| C7 | 31 | 54 | Ovary | Serous cystadenocarcinoma | T2N0M0 | II | malignant | 3+ | 3+ |
| C8 | 32 | 53 | Ovary | Serous cystadenocarcinoma | T1N0M0 | II~III | malignant | 3+ | 3+ |
| C9 | 33 | 22 | Ovary | Borderline mucinous cystadenoma | - | | borderline | Trace | Negative |
| C10 | 34 | 20 | Ovary | Mucinous cystadenocacinoma | T2N0M0 | | malignant | 3+ | 2+ |
| C11 | 35 | 47 | Ovary | Mucinous cystadenocacinoma | T1N0M0 | | malignant | 2+ | 1+ |
| C12 | 36 | 43 | Ovary | Mucinous cystadenocacinoma | T2N0M0 | III | malignant | 3+ | 3+ |
| D1 | 37 | 33 | Ovary | Serous cystadenocarcinoma | T1N0M0 | I~II | malignant | 1+ | 2+ |
| D2 | 38 | 34 | Ovary | Serous cystadenocarcinoma | T2N0M0 | II | malignant | 1+ | 2+ |
| D3 | 39 | 56 | Ovary | Serous cystadenocarcinoma | T1N0M0 | II | malignant | 1+ | 1+ |
| D4 | 40 | 45 | Ovary | Serous cystadenocarcinoma | T1N0M0 | II | malignant | 3+ | 2+ |
| D5 | 41 | 43 | Ovary | Serous cystadenocarcinoma | T1N0M0 | III | malignant | 1+ | 2+ |
| D6 | 42 | 51 | Ovary | Serous cystadenocarcinoma | T3N1M0 | III | malignant | 2+ | 2+ |
| D7 | 43 | 54 | Ovary | Serous cystadenocarcinoma | T2N0M0 | II | malignant | 3+ | 3+ |

| | | | | | | | | | |
|---|---|---|---|---|---|---|---|---|---|
| D8 | 44 | 53 | Ovary | Serous cystadenocarcinoma | T1N0M0 | II~III | malignant | 3+ | 3+ |
| D9 | 45 | 22 | Ovary | Borderline mucinous cystadenoma | - | | borderline | Trace | 1+ |
| D10 | 46 | 20 | Ovary | Mucinous cystadenocacinoma | T2N0M0 | | malignant | 1+ | 1+ |
| D11 | 47 | 47 | Ovary | Mucinous cystadenocacinoma | T1N0M0 | | malignant | Negative | 1+ |
| D12 | 48 | 43 | Ovary | Mucinous cystadenocacinoma | T2N0M0 | III | malignant | 3+ | 3+ |
| E1 | 49 | 18 | Ovary | Mucinous cystadenocacinoma | T1N0M0 | I | malignant | 2+ | 3+ |
| E2 | 50 | 25 | Ovary | Mucinous cystadenocacinoma | T1N0M0 | I~II | malignant | 1+ | 1+ |
| E3 | 51 | 24 | Ovary | Mucinous cystadenocacinoma | T1N0M0 | I | malignant | Negative | 1+ |
| E4 | 52 | 51 | Ovary | Endometrioid adenocarcinoma | T1N0M0 | I | malignant | 2+ | 3+ |
| E5 | 53 | 50 | Ovary | Endometrioid adenocarcinoma | T1N0M0 | I~II | malignant | 3+ | 3+ |
| E6 | 54 | 17 | Ovary | Endometrioid adenocarcinoma | T1N0M0 | I | malignant | 3+ | 3+ |
| E7 | 55 | 42 | Ovary | Endometrioid adenocarcinoma | T1N0M0 | I | malignant | 1+ | 2+ |
| E8 | 56 | 25 | Ovary | Endometrioid adenocarcinoma | T2N0M0 | I~II | malignant | 3+ | 3+ |
| E9 | 57 | 42 | Ovary | Endometrioid adenocarcinoma | T1N0M0 | II | malignant | 2+ | 3+ |

| | | | | | | | | | |
|---|---|---|---|---|---|---|---|---|---|
| E10 | 58 | 42 | Ovary | Endometrioid adenocarcinoma | T1N0M0 | II | malignant | 2+ | 3+ |
| E11 | 59 | 42 | Ovary | Endometrioid adenocarcinoma | T1N0M0 | II | malignant | 2+ | 3+ |
| E12 | 60 | 63 | Ovary | Endometrioid adenocarcinoma | T1N0M0 | III | malignant | 3+ | 3+ |
| F1 | 61 | 18 | Ovary | Mucinous cystadenocacinoma | T1N0M0 | I | malignant | 2+ | 3+ |
| F2 | 62 | 25 | Ovary | Mucinous cystadenocacinoma | T1N0M0 | I~II | malignant | 2+ | 3+ |
| F3 | 63 | 24 | Ovary | Mucinous cystadenocacinoma | T1N0M0 | I | malignant | 3+ | 1+ |
| F4 | 64 | 51 | Ovary | Endometrioid adenocarcinoma | T1N0M0 | I | malignant | 2+ | 3+ |
| F5 | 65 | 50 | Ovary | Endometrioid adenocarcinoma | T1N0M0 | I~II | malignant | 2+ | 3+ |
| F6 | 66 | 17 | Ovary | Endometrioid adenocarcinoma | T1N0M0 | I | malignant | 3+ | 3+ |
| F7 | 67 | 42 | Ovary | Endometrioid adenocarcinoma | T1N0M0 | I | malignant | 3+ | 2+ |
| F8 | 68 | 25 | Ovary | Endometrioid adenocarcinoma | T2N0M0 | I~II | malignant | 1+ | 3+ |
| F9 | 69 | 42 | Ovary | Endometrioid adenocarcinoma | T1N0M0 | II | malignant | 3+ | 2+ |
| F10 | 70 | 42 | Ovary | Endometrioid adenocarcinoma | T1N0M0 | II | malignant | 2+ | 2+ |
| F11 | 71 | 42 | Ovary | Endometrioid adenocarcinoma | T1N0M0 | II | malignant | 2+ | 3+ |
| F12 | 72 | 63 | Ovary | Endometrioid adenocarcinoma | T1N0M0 | III | malignant | 2+ | 3+ |
| G1 | 73 | 48 | Ovary | Endometrioid adenocarcinoma | T1N0M0 | III | malignant | 2+ | 3+ |

| | | | | | | | | | |
|---|---|---|---|---|---|---|---|---|---|
| G2 | 74 | 68 | Ovary | Endometrioid adenocarcinoma | T1N0M0 | III | malignant | 3+ | 3+ |
| G3 | 75 | 52 | Ovary | Endometrioid adenocarcinoma | T2N0M0 | III | malignant | 3+ | 3+ |
| G4 | 76 | 58 | Ovary | Endometrioid adenocarcinoma | T2N0M0 | III | malignant | 3+ | 3+ |
| G5 | 77 | 48 | Ovary | Undifferentiated carcinoma | T2N0M0 | III | malignant | 3+ | 3+ |
| G6 | 78 | 57 | Ovary | Undifferentiated carcinoma | T2N0M0 | III | malignant | 3+ | 2+ |
| G7 | 79 | 54 | Ovary | Undifferentiated carcinoma | T2N0M0 | III | malignant | 3+ | 2+ |
| G8 | 80 | 17 | Ovary | Dysgerminoma | T2N0M0 | | malignant | 3+ | 3+ |
| G9 | 81 | 40 | Ovary | Endodermal sinus tumor | T1N0M0 | | malignant | 1+ | 1+ |
| G10 | 82 | 34 | Ovary | Endodermal sinus tumor | T1N0M0 | | malignant | 2+ | 2+ |
| G11 | 83 | 36 | Ovary | Stromal sarcoma | T1N0M0 | | malignant | 3+ | 2+ |
| G12 | 84 | 31 | Ovary | Teratoma | | | benign | 2+ | 2+ |
| H1 | 85 | 48 | Ovary | Endometrioid adenocarcinoma | T1N0M0 | III | malignant | 3+ | 3+ |
| H2 | 86 | 68 | Ovary | Endometrioid adenocarcinoma | T1N0M0 | III | malignant | 3+ | 3+ |
| H3 | 87 | 52 | Ovary | Endometrioid adenocarcinoma | T2N0M0 | III | malignant | 3+ | 3+ |
| H4 | 88 | 58 | Ovary | Endometrioid adenocarcinoma | T2N0M0 | III | malignant | 3+ | 3+ |
| H5 | 89 | 48 | Ovary | Undifferentiated carcinoma | T2N0M0 | III | malignant | 3+ | 3+ |
| H6 | 90 | 57 | Ovary | Undifferentiated carcinoma | T2N0M0 | III | malignant | 3+ | 2+ |
| H7 | 91 | 54 | Ovary | Undifferentiated carcinoma | T2N0M0 | III | malignant | 3+ | 2+ |
| H8 | 92 | 17 | Ovary | Dysgerminoma | T2N0M0 | | malignant | 3+ | 3+ |
| H9 | 93 | 40 | Ovary | Endodermal sinus tumor | T1N0M0 | | malignant | 2+ | 1+ |
| H10 | 94 | 34 | Ovary | Endodermal sinus tumor | T1N0M0 | | malignant | 2+ | 1+ |
| H11 | 95 | 36 | Ovary | Stromal sarcoma | T1N0M0 | | malignant | 3+ | 2+ |

| | | | | | | | | | |
|---|---|---|---|---|---|---|---|---|---|
| H12 | 96 | 31 | Ovary | Teratoma | | | benign | 2+ | 1+ |

Remark: Detailed pathology report can be obtained online:
https://www.biomax.us/tissuearrays/Ovary/OV961

**Table S3: related to method section for RT² Profiler™ PCR array of human cell cycle genes illustrated in Fig 2.**

| No. | Genes | Description | $Log_2$ (Fold Change) | p-Value |
|---|---|---|---|---|
| 1 | ABL1 | C-abl oncogene 1, non-receptor tyrosine kinase | -1.994 | 0.000118 |
| 2 | ANAPC2 | Anaphase promoting complex subunit 2 | -2.2217 | 0.000017 |
| 3 | ATM | Ataxia telangiectasia mutated | -1.0346 | 0.810568 |
| 4 | ATR | Ataxia telangiectasia and Rad3 related | -2.3861 | 0.01008 |
| 5 | AURKA | Aurora kinase A | -1.7281 | 0.012692 |
| 6 | AURKB | Aurora kinase B | -2.0261 | 0.001432 |
| 7 | BCCIP | BRCA2 and CDKN1A interacting protein | -1.5577 | 0.167532 |
| 8 | BCL2 | B-cell CLL/lymphoma 2 | -1.8517 | 0.719269 |
| 9 | BIRC5 | Baculoviral IAP repeat containing 5 | -3.118 | 0.030112 |
| 10 | BRCA1 | Breast cancer 1, early onset | -1.6793 | 0.000205 |
| 11 | BRCA2 | Breast cancer 2, early onset | -2.3861 | 0.038622 |
| 12 | CASP3 | Caspase 3, apoptosis-related cysteine peptidase | 1.1423 | 0.387693 |
| 13 | CCNA2 | Cyclin A2 | -1.1662 | 0.087124 |
| 14 | CCNB1 | Cyclin B1 | -1.8615 | 0.049904 |
| 15 | CCNB2 | Cyclin B2 | -1.1754 | 0.113356 |
| 16 | CCNC | Cyclin C | -1.0992 | 0.441654 |
| 17 | CCND1 | Cyclin D1 | -3.6232 | 0.001663 |
| 18 | CCND2 | Cyclin D2 | 1.0851 | 0.70969 |
| 19 | CCND3 | Cyclin D3 | -2.8629 | 0.006304 |
| 20 | CCNE1 | Cyclin E1 | -2.9916 | 0.001867 |
| 21 | CCNF | Cyclin F | -2.1644 | 0.000135 |
| 22 | CCNG1 | Cyclin G1 | -1.618 | 0.024888 |
| 23 | CCNG2 | Cyclin G2 | -1.0851 | 0.937268 |
| 24 | CCNH | Cyclin H | -1.1053 | 0.430333 |
| 25 | CCNT1 | Cyclin T1 | -1.3084 | 0.088251 |

| | | | | |
|---|---|---|---|---|
| 26 | CDC16 | Cell division cycle 16 homolog (S. cerevisiae) | -1.197 | 0.094628 |
| 27 | CDC20 | Cell division cycle 20 homolog (S. cerevisiae) | -1.3008 | 0.034714 |
| 28 | CDC25A | Cell division cycle 25 homolog A (S. pombe) | -1.8996 | 0.007148 |
| 29 | CDC25C | Cell division cycle 25 homolog C (S. pombe) | -1.7838 | 0.008397 |
| 30 | CDC34 | Cell division cycle 34 homolog (S. cerevisiae) | -2.1737 | 0.000029 |
| 31 | CDC6 | Cell division cycle 6 homolog (S. cerevisiae) | -2.3341 | 0.002129 |
| 32 | CDK1 | Cyclin-dependent kinase 1 | -1.8172 | 0.027199 |
| 33 | CDK2 | Cyclin-dependent kinase 2 | -2.2964 | 0.001076 |
| 34 | CDK4 | Cyclin-dependent kinase 4 | -2.9036 | 0.001574 |
| 35 | CDK5R1 | Cyclin-dependent kinase 5, regulatory subunit 1 (p35) | -1.6599 | 0.020839 |
| 36 | CDK5RAP1 | CDK5 regulatory subunit associated protein 1 | -2.1744 | 0.01099 |
| 37 | CDK6 | Cyclin-dependent kinase 6 | -1.9379 | 0.007455 |
| 38 | CDK7 | Cyclin-dependent kinase 7 | -1.1013 | 0.344581 |
| 39 | CDK8 | Cyclin-dependent kinase 8 | -1.8608 | 0.004058 |
| 40 | CDKN1A | Cyclin-dependent kinase inhibitor 1A (p21, Cip1) | 1.3378 | 0.060831 |
| 41 | CDKN1B | Cyclin-dependent kinase inhibitor 1B (p27, Kip1) | -1.5694 | 0.00972 |
| 42 | CDKN2A | Cyclin-dependent kinase inhibitor 2A (melanoma, p16, inhibits CDK4) | 1.7934 | 0.339904 |
| 43 | CDKN2B | Cyclin-dependent kinase inhibitor 2B (p15, inhibits CDK4) | Not detected | 0.70969 |
| 44 | CDKN3 | Cyclin-dependent kinase inhibitor 3 | -1.9527 | 0.001263 |
| 45 | CHEK1 | CHK1 checkpoint homolog (S. pombe) | -1.7856 | 0.028136 |
| 46 | CHEK2 | CHK2 checkpoint homolog (S. pombe) | -1.1991 | 0.086923 |
| 47 | CKS1B | CDC28 protein kinase regulatory subunit 1B | -1.9398 | 0.002308 |
| 48 | CKS2 | CDC28 protein kinase regulatory subunit 2 | -1.1624 | 0.119128 |
| 49 | CUL1 | Cullin 1 | -1.7531 | 0.009382 |
| 50 | CUL2 | Cullin 2 | -1.1075 | 0.336613 |

| | | | | |
|---|---|---|---|---|
| 51 | CUL3 | Cullin 3 | -1.2301 | 0.085832 |
| 52 | E2F1 | E2F transcription factor 1 | -1 | 0.00015 |
| 53 | E2F4 | E2F transcription factor 4, p107/p130-binding | -2.2877 | 0.000123 |
| 54 | GADD45A | Growth arrest and DNA-damage-inducible, alpha | -2.3693 | 0.000357 |
| 55 | GTSE1 | G-2 and S-phase expressed 1 | -1.9723 | 0.000985 |
| 56 | HUS1 | HUS1 checkpoint homolog (S. pombe) | -1.8149 | 0.010749 |
| 57 | KNTC1 | Kinetochore associated 1 | -1.1817 | 0.141325 |
| 58 | KPNA2 | Karyopherin alpha 2 (RAG cohort 1, importin alpha 1) | -1.4221 | 0.00411 |
| 59 | MAD2L1 | MAD2 mitotic arrest deficient-like 1 (yeast) | -1.2159 | 0.04653 |
| 60 | MAD2L2 | MAD2 mitotic arrest deficient-like 2 (yeast) | -1.8777 | 0.00001 |
| 61 | MCM2 | Minichromosome maintenance complex component 2 | -2.247 | 0.000112 |
| 62 | MCM3 | Minichromosome maintenance complex component 3 | -1.9992 | 0.005942 |
| 63 | MCM4 | Minichromosome maintenance complex component 4 | -2.0199 | 0.000256 |
| 64 | MCM5 | Minichromosome maintenance complex component 5 | -2.6325 | 0.000357 |
| 65 | MDM2 | Mdm2 p53 binding protein homolog (mouse) | -2.4358 | 0.000199 |
| 66 | MKI67 | Antigen identified by monoclonal antibody Ki-67 | -1.6468 | 0.041091 |
| 67 | MNAT1 | Menage a trois homolog 1, cyclin H assembly factor (Xenopus laevis) | -1.6468 | 0.069017 |
| 68 | MRE11A | MRE11 meiotic recombination 11 homolog A (S. cerevisiae) | -1.3286 | 0.022893 |
| 69 | NBN | Nibrin | -1.3238 | 0.024218 |
| 70 | RAD1 | RAD1 homolog (S. pombe) | -2.5965 | 0.002259 |
| 71 | RAD17 | RAD17 homolog (S. pombe) | 1.0787 | 0.69002 |

| | | | | |
|---|---|---|---|---|
| 72 | RAD51 | RAD51 homolog (S. cerevisiae) | -1.8651 | 0.002835 |
| 73 | RAD9A | RAD9 homolog A (S. pombe) | -2.1558 | 0.00025 |
| 74 | RB1 | Retinoblastoma 1 | -1.7631 | 0.015369 |
| 75 | RBBP8 | Retinoblastoma binding protein 8 | -1.1536 | 0.220592 |
| 76 | RBL1 | Retinoblastoma-like 1 (p107) | -1.7685 | 0.005652 |
| 77 | RBL2 | Retinoblastoma-like 2 (p130) | -1.3298 | 0.140451 |
| 78 | SERTAD1 | SERTA domain containing 1 | -2.1528 | 0.000441 |
| 79 | SKP2 | S-phase kinase-associated protein 2 (p45) | -1.608 | 0.009959 |
| 80 | STMN1 | Stathmin 1 | -1.5416 | 0.060776 |
| 81 | TFDP1 | Transcription factor Dp-1 | -1.954 | 0.001492 |
| 82 | TFDP2 | Transcription factor Dp-2 (E2F dimerization partner 2) | -1.5096 | 0.025299 |
| 83 | TP53 | Tumor protein p53 | -1.3304 | 0.12448 |
| 84 | WEE1 | WEE1 homolog (S. pombe) | 1.1508 | 0.123016 |

**Table S4, related to results section for RPPA assay**

**List of genes selected for IPA with 0.4Log2 fold change illustrated in Fig 2.**

| No. | Genes | $Log_2$ (Fold Change) | p-Value |
|---|---|---|---|
| 1. | CDKN1A | 0.455304 | 7.80E-06 |
| 2. | HK2 | -0.40631 | 4.11E-05 |
| 3. | PRKAR1A | -0.33992 | 4.95E-05 |
| 4. | RRM2 | -0.36694 | 4.95E-05 |
| 5. | FOXO3 | -0.26697 | 4.95E-05 |
| 6. | TMEM173 | 0.408772 | 4.95E-05 |
| 7. | PAX8 | -0.35243 | 6.89E-05 |
| 8. | FGF2 | -0.3365 | 6.89E-05 |
| 9. | TRIP13 | -0.28622 | 6.93E-05 |
| 10. | DUSP4 | 0.259698 | 0.00010277 |
| 11. | PAR | 0.545277 | 0.00010277 |
| 12. | CDC6 | -0.41968 | 0.0001115 |
| 13. | MYC | -0.27618 | 0.00012527 |
| 14. | SGK1 | 0.248365 | 0.00012527 |
| 15. | KDR | 0.213099 | 0.00012527 |
| 16. | RAD23A | -0.35948 | 0.00017593 |
| 17. | FOXM1 | -0.14735 | 0.00017593 |
| 18. | EEF2K | -0.18532 | 0.00017593 |
| 19. | NOTCH3 | 0.192567 | 0.00017593 |
| 20. | RPS6KB1 | -0.33668 | 0.00019748 |
| 21. | MECOM | -0.2758 | 0.00019748 |
| 22. | ARHGAP45 | -0.18752 | 0.00020605 |
| 23. | DUSP6 | -0.08413 | 0.00027032 |
| 24. | SERPINE1 | 0.330056 | 0.00027032 |
| 25. | SGK3 | 0.205203 | 0.00028653 |
| 26. | CD276 | 0.374695 | 0.0005833 |
| 27. | CCNB1 | -0.24102 | 0.00063685 |
| 28. | RPS6KA1 | -0.12207 | 0.00063685 |
| 29. | CHEK1 | -0.26144 | 0.00094889 |
| 30. | AXL | 0.205664 | 0.00130672 |
| 31. | SLFN11 | -0.25799 | 0.00159804 |
| 32. | PTK2 | 0.21148 | 0.00348783 |
| 33. | ATM | -0.22364 | 0.012155 |

| | | | |
|---|---|---|---|
| 34. | ATP5F1A | -0.21571 | 0.01964946 |

**Table S5: related to method section of REMSA**

**Oligonucleotides used in REMSA experiment**

The oligonucleotides are derived from the region encompassing the AUUUA-rich regions in the human *cMYC* mRNA 3'UTR. The consensus sequences are indicated in boldface.

| Oligonucleotide | Sequence (5'-3') |
|---|---|
| 3'UTR-ARE1 | gaaag**auuua**gccau |
| 3'UTR-ARE2 | uuugu**auuua**agaau |
| 3'UTR-ARE3 | uuaag**auuua**cacaa |
| 3'UTR-ARE4 | uuuuu**auuua**aguac |
| 3'UTR-ARE5 | uugug**auuua**uuuug |
| 3'UTR-ARE6 | aacau**auuua**uucuu |
| 3'UTR-R1 | ucuguugaaaugggu |
| 3'UTR-R2 | gaaguagagagggaa |

**Table S6, related to method section of PLA**

**Oligonucleotides used in PLA experiment.**

The oligonucleotides are derived from the 7 nucleotides far from the ARE in the human *cMYC* mRNA 3'UTR.

| **Oligonucleotide** | **Sequence (5'-3')** |
|---|---|
| PLA probe1 | acugccucaaauuggacuuugggcauaaaa |
| PLA probe 2 | cuguaaauauugccauuaaauguaaauaac |
| PLA probe 3 | uuuuuaaaguugauuuuuuucuauuguuuu |
| PLA probe 4 | uuuguuguucauuguuuaauuuuuccuuuu |
| PLA probe 5 | ggagucauuauuuuagaaacuacaaacucu |

**Table S7, related to method section of luciferase reporter assay.**

**Oligonucleotides used in luciferase reporter experiment.**

| Oligo nucle otide | Sequence (5'-3') |
|---|---|
| 3'UTR | ggaaaaguaaggaaaacgauuccuucuaacagaaauguccugagcaaucaccuaugaacuuguuucaaaugcaugaucaaaugcaaccuc<br>acaaccuuggcugagucuugagacugaaagauuuagccauaauguaaacugccucaaauuggacuuugggcauaaaagaacuuuuuuaug<br>cuuaccaucuuuuuuuuucuuuaacagauuuguauuuaagaauuguuuuaaaaaauuuuaagauuuacacaauguuucucuguaaau<br>auugccauuaaauguaaauaacuuuaauaaaacguuuauagcaguuacacagaauuucaauccuaguauauaguaccuaguauuauagg<br>uacuauaaacccuaauuuuuuuuauuuaaguacauuuugcuuuuuaaaguugauuuuuuucuauuguuuuagaaaaaauaaaauaacu<br>ggcaaauauaucauugagccaaaucuuaaguugugaauguuuuguuucguuucuucccccucccaaccaccaccaucccuguuuguuuca<br>ucaauugccccuucagaggguggucuuaagaaaggcaagaguuuuccucuguugaaaugggucugggggccuuaaggucuuuaaguucuu<br>ggagguucuaagaugcuuccuggagacuaugauaacagccagaguugacaguuagaaggaauggcagaaggcaggugagaaggugagag<br>guaggcaaaggagauacaagaggucaaagguagcaguuaaguacacaaagaggcauaaggacuggggaguugggaggaaggugaggaag<br>aaacuccuguuacuuuaguuaaccagugccaguccccugcucacuccaaacccaggaauucugcccaguugaugggacacgguggaacc<br>agcuucugcugccuucacaaccaggcgccaguccuguccaugguuaucucgcaaaccccagaggaucucugggaggaaugcuacuauuaa<br>cccuauuucacaaacaaggaaauagaagagcucaaagagguuauguaacuuaucuguagccacgcagauaauacaaagcagcaaucugga<br>cccauucuguucaaaacacuuaacccuucgcuaucaugccuugguucaucugggucuaaugugcugagaucaagaagguuuaggaccuaa<br>uggacagacucaagucauaacaaugcuaagcucuauuugugucccaagcacuccuaagcauuuuaucccuaacucuacaucaaccccauga<br>aggagauacuguugauuuccccauauuagaaguagagagggaagcugaggcacacaaagacucauccacaugcccaagauucacugauag<br>ggaaaaguggaagcgagauuugaacccaggcuguuuacuccuaaccuguccaagccaccucucagacgacgguaggaaucagcuggcugcu<br>ugugaguacaggaguuacaguccagugguuauguuuuuaagucucaacaucuaagccuggucaggcaucaguuccccuuuuuuuguga<br>uuuauuuuguuuuuauuuuguuguucauuguuuaauuuuuccuuuuacaaugagaaggucaccaucuugacuccuaccuuagccauuugu<br>ugaaucagacucaugacggcuccugggaagaagccaguucagaucauaaaauaaaacauauuuauucuuugucaugggagucauuauuuu<br>agaaacuacaaacucuccuugcuuccauccuuuuuuacauacucaugacacaugcucauccugaguccuugaaagguauuuuugaacau<br>guguauuaauuauaagccucugaaaaccuauggcccaaaccagaaaugauguugauuauauagguaaaugaaggaugcuauugcuguuc<br>uaauuaccucauugucucagucucaaaguaggucuucagcucccuguacuuugggauuuuaaucuaccaccacccauaaaucaauaaaua<br>auuacuuucuuuga |

| | |
|---|---|
| Mut1 | ucuugagacugaaag**cgggc**gccauaauguaaacugc |
| Mut2 | Cuuuaacagauuugu**cgggc**agaauuguuuuuaaa |
| Mut3 | Acccuaauuuuuuuu**cgggc**aguacauuuugcuuu |
| Mut4 | Uccccuuuuuuugug**cgggc**uuuuguuuuuauuuu |

The oligonucleotides are derived from the region encompassing the AUUUA-rich regions in the human *cMYC* mRNA 3'UTR.

The base substitution mutant sequences of five nucleotides are indicated in boldface.
